# Machine learning-guided optimization of p-coumaric acid production in yeast

**DOI:** 10.1101/2023.11.27.568789

**Authors:** Sara Moreno-Paz, Rianne van der Hoek, Elif Eliana, Priscilla Zwartjens, Silvia Gosiewska, Vitor A.P. Martins dos Santos, Joep Schmitz, Maria Suárez-Diez

**Affiliations:** Wagenigen University and Research; DSM-firmenich; Wageningen University and Research; Wageningen University & Research

## Abstract

Industrial biotechnology uses Design-Build-Test-Learn (DBTL) cycles to accelerate the development of microbial cell factories, required for the transition to a bio-based economy. To use them effectively, appropriate connections between each phase of the cycle are crucial. Using p-coumaric acid production in *Saccharomyces cerevisiea* as case study, we propose the use of one-pot library generation, random screening, targeted sequencing and machine learning (ML) as links during DBTL cycles. We showed that the robustness and flexibility of ML models strongly enable pathway optimization, and propose feature importance and SHAP values as a guide to expand the design space of original libraries. This approach led to a 68% increased production of p-coumaric acid within two DBTL cycles.

## 1 Introduction

The reality of climate change calls from an imminent transition to a bio-based economy less reliable on the petrochemical industry. Biotechnology contributes to solutions to this problem as metabolic engineering allows microbial production of a wide variety of compounds such as pharmaceuticals, biofuels, food additives and bulk chemicals [1]. However, these solutions often require very long development times that limit their real-world application [2].

Design-Build-Test-Learn (DBTL) cycles offer a framework for systematic metabolic engineering. Pathways are designed during the Design phase, strains are constructed in the Build phase and screened for production during the Test phase. In the Learning phase, a relationship between pathway design and production is established and used to inform new DBTL cycles [3]. Advances on synthetic biology and automation facilitate the engineering of microorganisms and increase the throughput of the Build and Test phases. However, predicting the effect of modifications in the Design phase that may lead to improvements is non-trivial [4, 5]. In fact, the acceleration of the Build and Test phases of the DBTL cycle might lead to a paradox where more data leads to more complexity but not necessarily better strain performance [6]. To avoid this, an efficient, meaningful link between the design and learn phases of the cycle is crucial.

Machine learning (ML) can identify patterns in the system of interest without the need of detailed mechanistic understanding of the problem [7]. It has been used to aid strain development with applications ranging from gene annotation and pathway design to process scale-up [4]. When used for pathway optimization, common approaches start by creating libraries of strains with varying regulatory elements (e.g. promoters, ribosome binding sites). These libraries include a defined solution space that can be explored by random or rational sampling [8, 9]. A subset of the library is then screened, and genotype and production data are used to train ML algorithms. The algorithms then suggest a new round of (improved) strains for construction, effectively linking the Learn and Design phases of sequential DBTL cycles [5, 8, 10–12]. Besides, ML algorithms are robust to missing data caused by unsuccessful construction of specific strains which facilitates effective and efficient implementation of DBTL cycles [12, 13].

p-Coumaric acid (pCA) is an aromatic amino acid-derived molecule produced from phenylananine (Phe) or tyrosine (Tyr). It is naturally found in plants and serves as starting material for commercially valuable products such as pharmaceuticals, flavors, fragrances, and cosmetics [14]. In *Saccharomyces cerevisiae* Phe and Tyr are synthetized via the phrephenate pathway (Figure 1A) [15, 16]. This pathway starts with the condensation of erythrose-4-phosphate (E4P) and phosphoenolpyruvate (PEP) by 3-deoxy-7-phosphoheptulonate synthase (ARO3/4). Then, the pentafuntional protein ARO1, converts 3-deoxy-7-phosphoheptulonate (DAHP) to 5-enolpyruvylshikimate-3-phosphate (EPS3P), that is converted to chorismate (CHO) by ARO2, and to phrephenate (PRP) by ARO7. Phrephenate can then be converted to phenylalanine by prephenate dehydratase (PHEA) and ARO8/9, or to tyrosine by prephenate dehydrogenase (TYR) and ARO8/9. To continue the synthesis of pCA, expression of heterologous genes is needed: tyrosine ammonia lyase (TAL) for synthesis from Tyr; or phenylalanine ammonia lyase (PAL), cinnamate 4-hydroxylase (C4H) and its associated cytochrome P450 reductase (CPR) for synthesis from Phe [14, 17–19].

**Figure 1:**
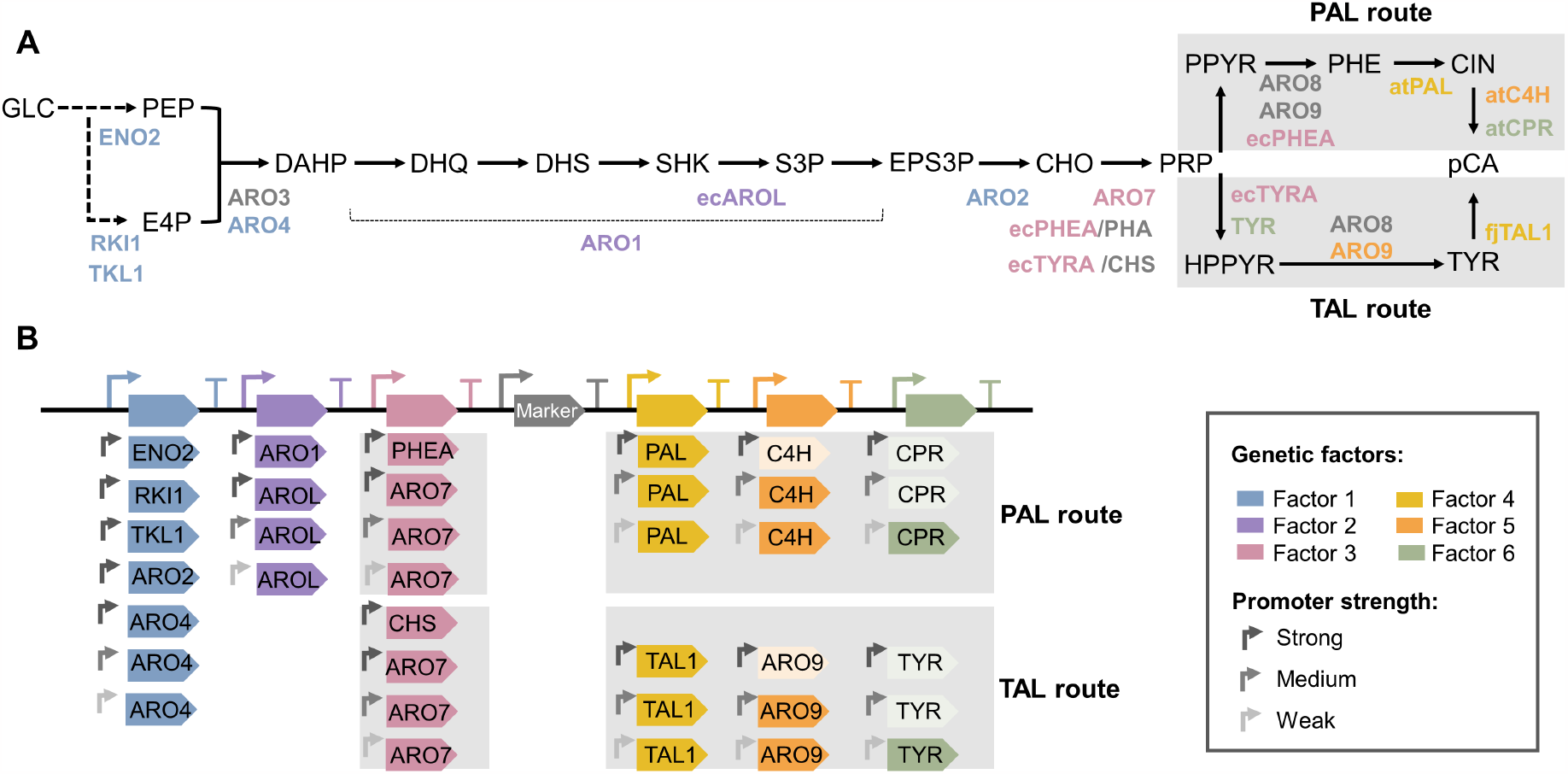
**A**. pCA production pathway. Heterologous genes are preceded by a two-letter code indicating the organism of origin where ”ec” refers to *E. coli*, ”at” to *Arabidopsis thaliana* and ”fj” to *Flavobacterium johnsoniae*, see legend for colour codes . **B**. Library structure. The library consists of gene clusters formed by a selection marker (Marker) and six factors with levels including different open reading frames (ORFs) and different promoters. Lighter colors are used to indicate factor’s levels included in the design but not obtained experimentally. GLC, glucose; PEP, phosphoenolpyruvate; E4P, erithrose-4-phosphate; DAHP, 3-deoxy-7-phosphoheptulonate; DHQ, 3-dehydroquinate; DHS, 3-dehydroshikimate; SHK, shikimate; S3P, shikimate-3-phosphate; EPS3P, 5-enolpyruvylshikimate-3phosphate; CHO, chorismate; PRP, phrephenate; PPYR, phenylpyruvate; PHE, phenylalanine, CIN, cinnamate; pCA, p-coumaric acid; HPPYR, 4-Hydroxyphenylpyruvate; TYR, tyrosine.

The prephenate pathway is highly regulated where Tyr exerts feedback inhibition on ARO3 and ARO7 and Phe on ARO4 [16]. This regulation together with availability of precursors and appropriate expression of heterologous genes have demonstrated to influence pCA production [14, 17, 18]. However, testing the effect of these factors individually might result in the exclusion of possible synergistic effects. Alternatively, combinatorial optimization of metabolic pathways can facilitate the search of optimal production albeit involving the construction and testing of an exponentially growing number of strains [8].

We used ML-guided DBTL cycles to improve pCA production in *S. cerevisiae*. We created combinatorial libraries based on the Tyr or Phe-derived pathways, that simultaneously altered expressed coding sequences and regulatory elements (promoters) (Figure 1B). We showed a better performance of the Phe-derived pathway, which was further optimized based on ML predictions. Following this strategy we achieved a 68% improvement in production within two DBTL cycles and a final pCA titer of 0.52 g/l .

## 2. Materials and methods

### 2.1 Organisms and media

*S. cerevisiae* strains were derived from CEN.PK113-7D and grown at 30°C in Yeast Extract Phytone Dextrose media (YEPhD, 2% Difco™ phytone peptone (Becton-Dickinson (BD), Franklin Lakes, NJ, USA), 1% Bacto™ Yeast extract (BD) and 2% D-glucose (Sigma Aldrich, St Louis, MO, USA)). When required, antibiotics were added to the media at appropriate concentrations: 200 µg/ml nourseothricin (Jena Bioscience, Germany), 200 µg/ml geneticin (G418, Sigma-Aldrich). *E. coli* DH10B (New England BioLabs, Ipswich, MA, USA) was used as cloning strain and grown at 37°C in 2*Peptone Yeast Extract mdia (2*PY, 1.6% tryptone peptone (BD), 1% Bacto™ yeast extract (BD) and 0.5% NaCl (Sigma Aldrich)). When required, antibiotics were added to the media at appropriate concentrations: 100 µg/ml ampicillin (Sigma-Aldrich), 50 µg/ml neomycin (Sigma-Aldrich). Solid medium was prepared by addition of Difco™ granulated agar (BD) to the medium to a final concentration of 2% (w/v).

### 2.2. Cassette construction

DNA templates for promoters and terminators [20] as well as open reading frames (ORF) were codon optimized [21] and can be found in Sup. Table 1. Bricks were assembled into cassettes (promoter + ORF + terminator) via Golden Gate (using BsaI-HF v2.0 (NEB) and T4 DNA Ligase (Invitrogen)) into a backbone plasmid containing a 50bp homologous connector sequence to facilitate in vivo recombination of the gene cluster as described in [22] (Sup. Figure 1). Golden Gate products were transformed into chemically competent *E. coli* DH10B. Wizard® SV 96 Plasmid DNA purification system (Promega, Madison, WI, USA) was used for plasmid isolation. Cassettes were confirmed by PCR using Q5® High-Fidelity DNA polymerase (NEB) with primers from IDT (Leuven, Belgium) and analyzed on a LabChip® GX Touch Nucleic Acid Analyzer (Perkin Elmer). Plasmids with correct fragment size were amplified by PCR using Q5® High-Fidelity DNA polymerase (NEB) and integration site flanks (50 base pair homologous region) were attached to the first and the last cassette of the gene cluster (Sup. Figure 1). PCR products were purified using Promega Wizard® SV PCR Clean-Up kit and quantified using DropSense 96 (Trinean).

### 2.3. Strain construction

Strains were constructed as described in [22, 23]. In short, host strain SHK001 pre-expressing Cas9 (integrated on locus INT1, Sup. Table 1) was used to enable a targeted integration via CRISPR-Cas9 into *S. cerevisiae*’s genome. A linear guide RNA targeting a single locus (Sup. Table 5) was amplified from a gBlock (IDT) with 50bp homology regions to pRN1120. Plasmid backbone pRN1120 was amplified for *in vivo* assembly of the gRNA plasmid. PCRs were performed with Q5 ® High-Fidelity DNA polymerase (NEB). PCR products were confirmed on a 0.8% agarose gel and purified using Wizard® SV Gel and PCR Clean-Up kit (Promega). DNA fragments were quantified using Nanodrop (ThermoFisher Scientific). Primers used are provided in Sup. Table 4.

Equimolar amounts (100-300 ng/kb) of the cassettes, linear gRNA (210 ng/kb) and linear backbone (35 ng/kb) fragments were transformed to the cells following the LiAc/ssDNA/PEG method [24]. Reagents required for yeast transformation were obtained from Sigma-Aldrich (lithium acetate dihydrate (LiAc) and deoxyribonucleic acid sodium salt from salmon testes (ssDNA)) and Merck (polyethylene glycol 4000 (PEG)). The connector sequences on the cassettes facilitates *in vivo* recombination of a cluster of genes in the genome [22]. Transformants were plated on Qtray (NUNC) containing YEPhD agar medium and selection agent. Colonies appeared on plate after 3 days incubation at 30°C. Single colonies were picked with Qpix 420 (Molecular Devices) into 96 well plate containing YEPhD agar medium and selection agent and regrown for 3 days at 30°C.

### 2.4 Whole genome sequencing

*S. cerevisiae* cells (OD 5-10) were pelleted and lysed in 200 µL 0.9% physiologic salt supplied with 2 µL RNAse cocktail (Invitrogen) and 5mg/mL Zymolyase 100T (MP Biomedicals). The mixture was incubated at 37°C for 45 minutes. Next, 200 µL 2X cell lysis solution (0.05M EDTA, 4%SDS) was added to the mixture, followed by vortexing. 168 µL protein precipitation solution (10M NH4Ac) was added and proteins were precipitated by centrifugation for 10 minutes at 20K rcf at 4°C. The DNA in the supernatant was precipitated with an equal volume of isopropanol (centrifugation for 2 minutes at 16K rcf at room temperature). The DNA pellet was washed with 70% ethanol. The ethanol was discarded and the pellet was left to dry and then dissolved in MilliQ water. The isolated genomic DNA was quantified using Qubit (Thermofisher Scientific) and Nanodrop (Thermofisher Scientific), purified using Zymo Research gDNA Clean & Concentrator kit and sequenced using the ligation sequencing kit (LSK-SQK109) with the native barcoding expansion (EXP-NBD114) from Oxford Nanopore Technologies according to manufacturer instructions on a GridION device (FLOW-MIN106 flow cell).

### 2.5 Promoter-terminator characterization

Combinations of promoter-terminator were characterized using GFP as reporter gene (see Sup. Table 3 for details). Precultures were prepared in 96-well half-deep well plates (HDWP) containing 350 µL YEPhD + Pen/Strep (Invitrogen) and incubated at 30°C, 750 rpm, 80% humidity for 48 hours. 10 µL of the grown pre-culture was re-inoculated to MTP-R48-B FlowerPlate (m2p-labs) containing 1 mL minimal medium + Pen/Strep (Invitrogen). The plate was incubated 48 h in Biolector® at 30°C, 800 rpm, 85% humidity. Biomass (em. 620nm/ex.620nm) and fluorescence (em.488nm/ex. 520nm), each with 3 filters (gain of 100, 50, and 20), were measured every 15 minutes. 40 µL of 2 days-old main culture were measured using fluorescence-activated cell sorting (BD, FACSAria Fusion) to detect single cells expressing GFP at flow rate of 10.000 evt/s. The signal of fluorescent proteins was detected with a bandpass filter set at 530/30 nm for eGFP. The data was recorded using BD FACSDiva 8.0.2 software to retrieve the geometric mean of the fluorescence distribution. Data was analyzed using FlowJo (version 10.6.2).

### 2.6 p-Coumaric acid production experiments

Colonies were grown in 96 microtiter plate (MTP) Nunc flat bottom (ThermoFisher Scientific) containing YEPhD and appropriate selection agent for 48 h at 30°C, 750 rpm, 80% humidity. Cultures were re-inoculated in HDWP (ThermoFisher Scientific, AB-1277) containing 350 µL YEPhD and selection agent and grown for 48 h in the same conditions. The grown cultures were reinoculated to HDWP containing 350 µL minimal media (Verduyn Luttik with 2% glucose [25]) and incubated for 2 days at 30°C, 750 rpm, 80% humidity. In all plates blank wells and wells containing a control strain (SHK0046, see Sup. Table 3) were included. For flow-NMR measurements 250 µL broth were sampled to a 96-deep well plate (DWP) and mixed with 500 µL Acetonitrile (Sigma Aldrich) by pipetting. The mixture was centrifuged at 4000 rpm for 10 minutes. 500 µL supernatant was transferred to a new DWP for analysis with flow-NMR. For LC/MS measurements 250 µl broth was sampled. 1 ml acetonitrile was added, sample was mixed by pipetting and centrifuged. 250 µl supernatant was diluted with 375 µl milliQ and used for analysis with LC/MS.

### 2.7 p-Coumaric acid quantification with automated segmented-flow NMR analysis

The DWP plates were lyophilized to remove the non-deuterated solvents. 100 µl solution of 1 g/l internal standard 1,1-difluoro-1-trimethylsilanyl methylphosphoric acid (FSP, Bridge Organics) in MilliQ water was added into DWP prior the lyophilization. To the lyophilized samples, 600 µl of D_2_O (Cambridge Isotope Laboratories (DLM-4)) was added and homogenized. The samples were analyzed on a CTC PAL3 Dual-Head Robot RTC/RSI 160 cm robotic autosampler (CTC Analytics AG, Zwingen, Switzerland) fluidically coupled to a Bruker spectrometer Avance III HD 500 MHz UltraShield [26]. ^1^H spectra were recorded with standard pulse program (zgcppr) with following parameters: 16 scans, 2 dummy scans, 33k data points, 16.4 ppm spectral width, 1.2 s relaxation delay (d1), 8 µs 90° pulse, 2s acquisition time, 15 Hz water suppression, and fixed receiver gain (rg) of 64 Spectra were processed and analyzed using Topspin 4.1.4 (Bruker). Spectral phasing was applied and spectra were aligned to 3-(trimethylsilyl)-1-propanesulfonic acid-d6 sodium salt (DSS-d6, Sigma-Aldrich) at 0 ppm. Auto baseline correction was applied on the full spectrum width. Additional third-order polynomial baseline correction for selected regions was applied if needed. The amount of pCA (doublet, 6.38 ppm, n=2H) was calculated relative to the signal of FSP. NMR production data per plate was normalized by SHK0046 production.

### 2.8 p-Coumaric acid quantification with LC-HR-MS spectrometry

Samples were analyzed on an Vanquish Horizon UHPLC system coupled to a Q Exactive Focus mass spectrometer (ThermoFisher). The chromatographic separation was achieved on a Acquity UPLC® BEH C18 column (100 x 2.1 mm, 1.7 µm, Waters), using gradient elution with (A) 0.025% formic acid in LC-MS grade water, and B) 90% LC-MS grade acetonitrile (Sigma Aldrich) with 10% mobile phase A with run time of 9 min. The gradient started with 1% B linearly increased to 50% B in 5 min, followed by rapid increase to 99% B in 0.1 min and, kept at 99% for 1.9 min and then re-equilibrated with 1% B for 1.9min. The flow rate was kept at 0.6 ml/min, using an injection volume of 2 µl and the column temperature was set to 50°C. pCA was detected in negative APCI mode using and quantified using an external calibration line of a reference standard. Using this chromatographic system, the coumaric acid elutes at retention times 3.05 minutes with *m/z* 163.0403 (M-H), in good agreement (within 2 ppm) with the theoretical m/z value of 163.04007.

### 2.9 Machine learning-guided strain design

440 randomly screened strains were classified into four clusters based on NMR pCA production data. Strains from every cluster were randomly selected for sequencing in order to cover the complete solution space. Colonies were considered correct when they had targeted integration of the complete gene cluster (7 cassettes, one per factor and the selection marker). Genoptype and production data from correct colonies were used as input for modeling. For gene clusters present in more than one correct sequenced colony, average pCA production was considered. Two datasets were used: a *complete dataset* including data from producers and non-producers and a *producers dataset*. Colonies were considered non-producers when measured pCA production was below 0.05 a.u.

The following regressor models from scikit-learn were considered for evaluation: multiple linear regression (MLR), support vector regression (SVR), random forest regression (RFR) and kernel ridge regression (KRR). pCA production was modeled as a function of the different factor levels (genes or their expression strength), treated as categorical variables using one-hot-encoding. Each dataset was split into train (90% data) and test sets (10%) using stratification (*i*.*e*. maintaining the proportion of the different classes in both sets). For all models except MLR, hyper-parameters were selected based on leave-one-out cross-validation in the train set using the maximum error as score. Predictions of models with optimized hyper-parameters were compared to the test set using the coefficient of determination (R^2^) as score. This process was repeated ten times and models were compared based on their average R^2^ on the test sets (Sup Figure 4). Additionally, the impact of the training data size on model performance was tested: after the train test split, percentages of the training data from 5 to 100% were used for training and model performance was evaluated using the test set with R^2^ as score.

For each dataset, models selected based on R^2^ were trained following two different strategies: “*one time training*” and “*recurrent training*”. In the first strategy, all data from the dataset was used for model training. In the second strategy, 90% of the data from the dataset was used for model training and this process was repeated 100 times. Trained models were used to predict pCA production for all the designs in the design space. For each dataset and training strategy, top producers were ranked based on the frequency of each design being predicted as top 1, top 5 and top 10 by each model (Sup Figure 4).

The impact of the different factors on pCA production was evaluated by permutation feature importance using the permutation importance function from the inspection module of the scikit-learn library. In addition, SHapley Additive exPlanations (SHAP) values were calculated using the shap library [27].

All data and scripts used are available in [GitLab](https://gitlab.com/wurssb/Modelling/ml4pca). Model selection, training and feature importance was performed using Python 3.8.8 and Scikit-learn 1.1.3 [28].

## 3 Results

### 3.1 DBTL Cycle 1: Exploring the design space

#### 3.1.1 Design: selection of factors and levels

Two independent libraries were designed depending on whether pCA was produced from Phe (PAL route) or Tyr (TAL route) (Figure 1A). Any design of the libraries is formed by a 7-genes cluster (6 factors and a selection marker) integrated in the genome of *S. cerevisiae* (Figure 1B, Sup Figure 1). The combination of promoter, ORF and terminator (cassette) in the gene cluster constitutes a factor that can take different levels depending on the chosen promoter and/or ORF. The size of the library is determined by the number of factors and levels so 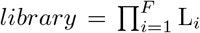, where *F* is the number of factors and *L*_*i*_ is the number of levels of factor *i*. Both libraries shared factors 1 and 2 and differed in the other 4 factors (Figure 1B).

Factor 1 contained five ORFs: enolase (ENO1), ribose-5-phosphate isomerase (RKI1), transketolase (TKL1), ARO2 and a feedback-resistant ARO4 (ARO4^K229L^) under the TDH3 promoter. Besides, ARO4^K229L^ could be downstream of two additional promoters (RPL8A, MYO4) as expression of this gene has resulted in significantly increased pCA titers [14, 18]. ENO1, RKI1 and TKL1 were chosen considering that availability of PEP and E4P can also affect production. ARO2 was included as additional level to test the effect of other shikimate pathway genes.

Levels for factor 2 were based on the assumption of ARO1 as rate limiting step. Rodriguez et al. observed increased pCA production when ARO1 or AROL from *E. coli*, which catalyzes the phosphorylation of shikimate, were over-expressed in yeast [18]. Therefore, 4 levels were chosen: expression of AROL under three different promoters (TEF1, RPL28, UREA3) and expression of ARO1 under a strong promoter (TEF1).

The focus of factor 3 was on the expression of the feedback-resistant variant ARO7^G141S^ under three different promoters (PRE3, ACT1, PFY1), as expression of this gene improved pCA titers [14, 18]. Besides, expression of PHEA and TYRA from *E. coli* with the PRE3 promoter are considered as additional levels for the PAL and TAL libraries respectively. These bifuncional enzymes have a chorismate mutase activity (conversion of CHO to PRP) and either phrephenate dehydratse (PHEA) or dehydrogenase (TYRA) activity, specific for the formation of Phe or Tyr respectively [29, 30].

Factors 4, 5 and 6 of the PAL library each focused on one of the heterologous genes required for pCA production from Phe: PAL, C4H and CPR under the control of three different promoters (ENO2, RPS9A, VMA6; KI OLE1, CHOI, PXR1; and PGK1, RPS3, CCW12 respectively). In the TAL library, levels of factor 4 were formed by TAL under the control of three promoters (ENO2, RPS9A, and VMA6). In order to obtain a design space with the same size as the PAL library, factors 5 and 6 included the expression of ARO9 and TYR with the same promoters used for the PAL library.

Considering the factors and levels used, the number of possible designs in each library was 3024 (7 *·* 4 *·* 4 *·* 3 *·* 3 *·* 3).

#### 3.1.2 Build and Test: construction and screening of the combinatorial library

For each of the promoter-terminator pairs designed, cassettes formed by promoter-GFP-terminator were constructed and transformed into yeast. Positive colonies were found for all the constructs but the strong promoter-terminator pairs for factors 3, 5 and 6 (PRE3-ADH1, OLE1-TDH3, and PGK1GPM1). Cells were grown in BioLector bioreactors and fluorescence was analyzed using FACS. For factor 1, fluorescence of strong and medium promoters differed by an order of magnitude. For factors 2 and 4, fluorescence values for the medium promoters were approximately half of those from strong promoters. Weak promoters showed fluorescence values 1 or 2 order of magnitudes below the strong and medium promoters (Sup. Figure 2).

Cassettes required for the *in vivo* assembly of the gene clusters were created combining promoters, ORFs, terminators and homology regions. All cassettes except those containing the strong promoter for factor 5 and the strong and medium promoters for factor 6 were obtained, which reduced the size of the PAL and TAL libraries from 3024 possible designs to 672 (Figure 1B).

*S. cerevisiae* cells expressing Cas9 were transformed with a mixture of the correct cassettes using onepot transformation. Cells were plated in selective media and 440 strains per library were randomly selected for screening of pCA production. These stains were grown in 96DWP for 48h, pCA was extracted and samples were measured using NMR (Figure 2). Colonies from the PAL route produced pCA ranging from 0 to 0.22 a.u. Colonies from the TAL route produced significantly less pCA, with only three colonies producing above the detection limit (0.05 a.u.) and a maximum production of 0.10 a.u.

**Figure 2:**
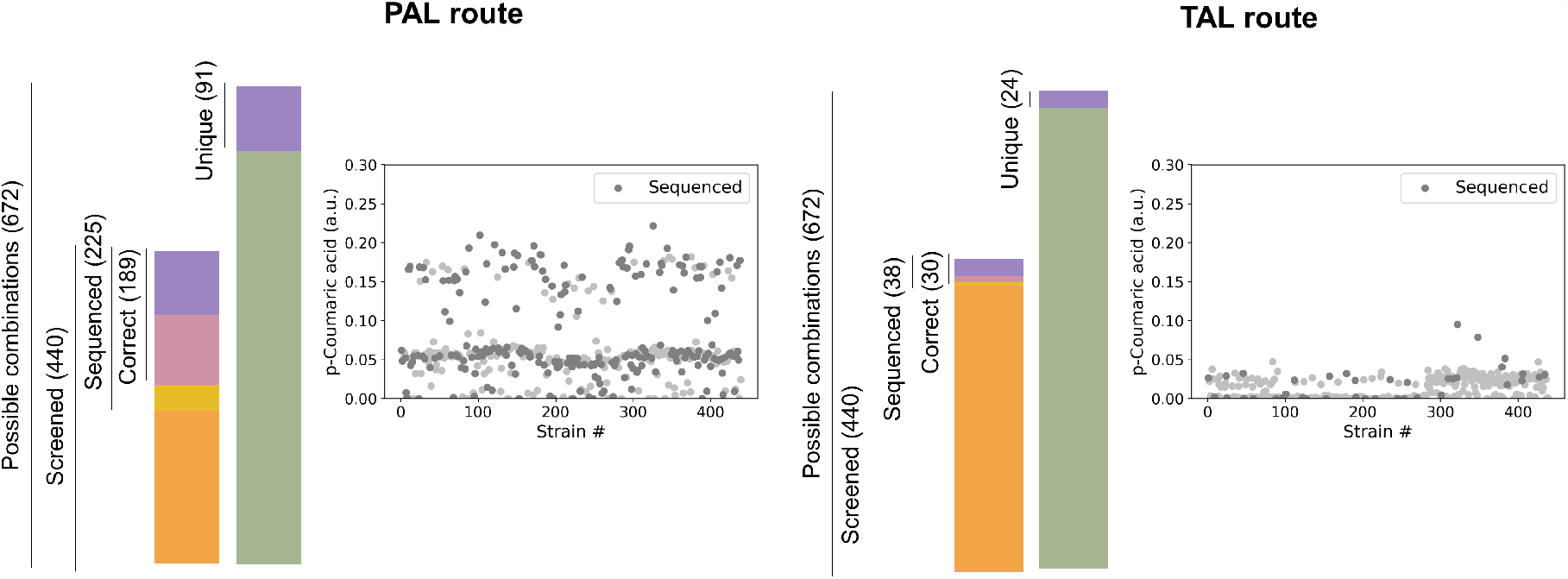
Screening before sequencing strategy. For each of the routes allowing pCA production, 672-member libraries were defined. For the PAL route, production of 440 randomly selected strain was measured and 225 strains were selected for sequencing; 189 correct strains containing 91 unique pathway designs were found. For the TAL route, 440 strains were screened from which 38 were sequenced; 30 of these strains were correct and 24 unique designs were found.

Considering the screening results, the production space was sampled including low, medium and high producers in order to obtain high-quality data for ML [4] and to analyze the efficiency of the library generation method. For the TAL route, 38 strains were sequenced from which 30 sequences were correct (*i*.*e*. contained the gene cluster with 7 genes) and 24 contained unique pathway designs(Figure 2). Considering that 80% of the sequenced strains were correct, the observed low pCA production was likely caused by the lower efficiency of the TAL route and not incorrect library construction. These results agreed with previous reports that identified the PAL route as the most suitable pathway for pCA production [14]. Therefore, the optimization of pCA production was focused on the PAL route, from which 225 strains were sequenced. We found 189 correct strains (84%) from which 91 (48%) were unique, validating the library construction approach (Figure 2). Out of the 91 unique designs, 58 designs were present in one strain and 33 had multiple replicates (Sup. Figure 3). Besides, for all factors at least a strain containing each of the levels was found (Figure 3A).

**Figure 3:**
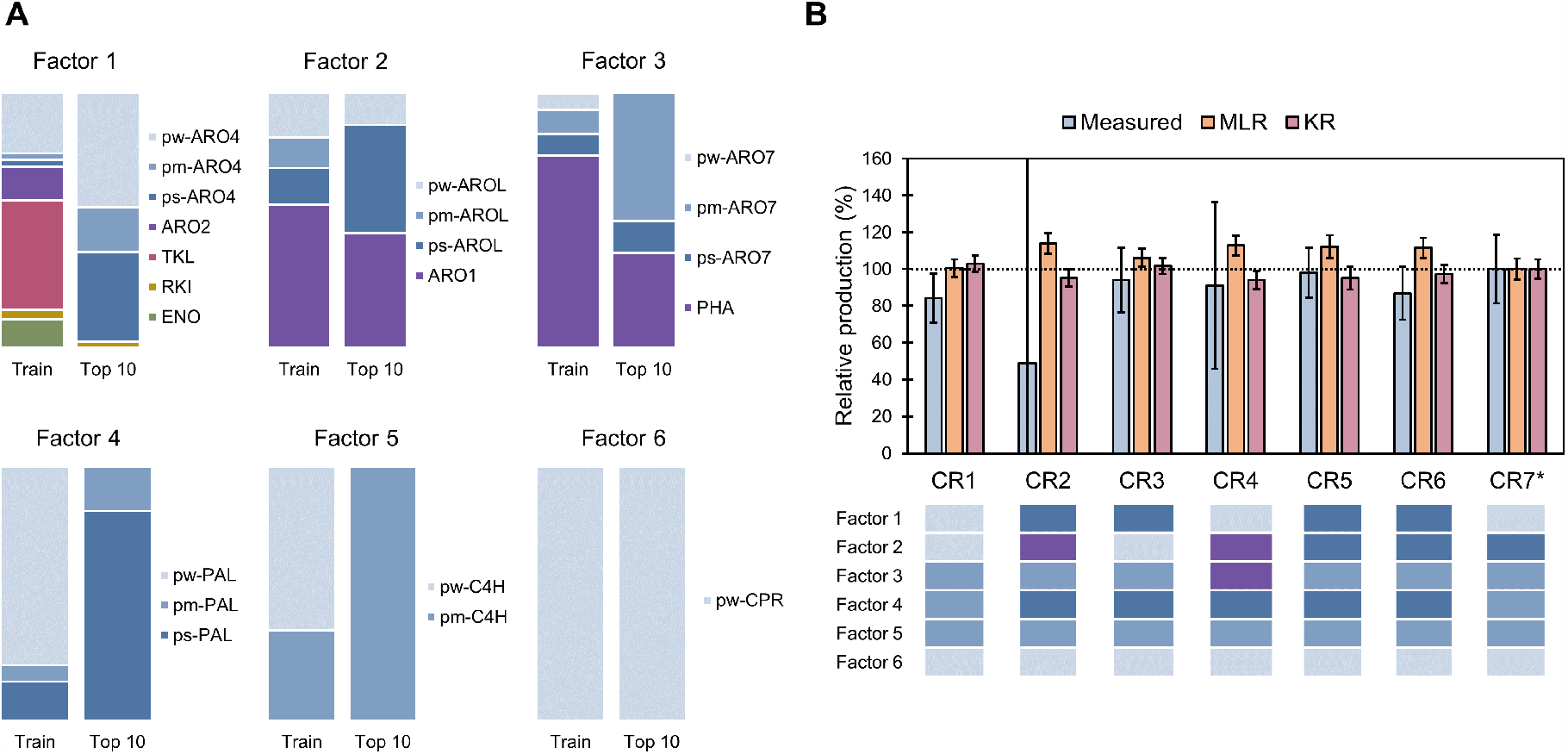
**A**. Comparison of factor’s levels on training set and top 10 predicted strains considering the CO, CR, PO and PR rankings. Ps, pm and pw indicate strong, medium and weak promoters respectively. **B**. Experimental validation of the CR ranking predictions. Production relative to the BMP strain (same as CR7* strain) is shown. Genotype of the strains follows same colour code as panel A.

#### 3.1.3 Learn: model selection, training and predictions

One of the challenges of applying ML to strain design is the training data requirements. While some reports suggest the homogeneous sampling of the complete solution space [4], other suggest the benefit of including mainly good producers [10]. Therefore, we divided our data in two datasets: the *complete* dataset that included data from producers and non-producers and the *producers* dataset. Stratification was used during training to ensure constant proportion of poor, medium, good and very good producers in the train and test sets. Train sets were used to find optimal hyper-parameters for four ML algorithms: MLR, SVR, KRR and RFR. Performance of the models with optimized hyperparameters was evaluated on the test set (Sup Figure 4). Models trained with the producers dataset showed better performance than those trained using the complete dataset (Table 1). MLR and KRR or all models were chose as predictors for the complete and the producer datasets, respectively.

**Table 1:** Performance of ML methods (R^2^) trained with the complete or producer datasets using stratification. MLR, multiple linear regression; SVR, support vector regression; KRR, kernel ridge regression; RFR random forest regression.

|  | **Complete dataset**<br>(91 designs) | **Producers dataset**<br>(63 designs) |
| --- | --- | --- |
| **MLR** | $0.70 \pm 0.17$ | $0.82 \pm 0.15$ |
| **SVR** | $0.71 \pm 0.23$ | $0.82 \pm 0.19$ |
| **KRR** | $0.72 \pm 0.18$ | $0.80 \pm 0.11$ |
| **RFR** | $0.72 \pm 0.19$ | $0.82 \pm 0.16$ |

Selected models were trained in each dataset using two different learning strategies: “*one-time training*” and “*recurrent training*” (Sup Figure 3). The first strategy consisted of one-time training with all the available data and did not provide uncertainty in the predictions. The second strategy was based on recurrent learning on 90% of the available data which reduced the impact of possible outliers in the training data and allowed uncertainty quantification of predictions. Trained models were then used to predict pCA production of the 672 designs from the full design space. Considering that models selected for each dataset had similar performances (Table 1), designs were ranked based on the frequency in which each design was predicted to be in the top 1, top 5 or top 10 by each model. In this way the construction of designs commonly predicted as top producers by different models was favoured. Four rankings were obtained: the CO and CR rankings based on the Complete dataset and the One-time or Recurrent training strategies respectively, and the PO and PR rankings based on the Producers dataset (Sup Figure 4).

Training strategies were evaluated based on their ranking of the best measured producer strain (BMP), the five best measured producers (5-BMPs), and all the measured non-producers (Sup Figure 5). Best measured producers were expected to rank high while measured non-producers were expected to hold lower positions. Regardless of the training strategy, including non-producers during training did not change predictions of measured top producers, but improved predictions of measured non-producers, ensuring correct coverage of the complete design space by the ML predictions.

In order to improve pCA production, designs predicted to render the highest titers were evaluated (Sup Figure 6). Notably, the BMP strain was predicted as part of the top 10 designs in all but the CO ranking. A comparison between the levels present in the training data and in the top 10 designs predicted by all the learning strategies is depicted in Figure 3A and Sup Figure 6. Top predicted strains showed a preference for ARO4 under weak or strong promoters compared to the other ORFs. For factors 2 and 3, ARO1 or AROL under a strong promoter and PHA or ARO7 under its medium promoter were favoured. Finally, the stronger promoters tested for PAL and C4H were enriched in the predicted top producers.

However, predicted pCA production improvements compared to the BMP strain were low (6 *±* 8%, 2*±* 5%, 3 *±* 6% and 1 *±* 5% depending on the learning strategy used) (Sup Figure 7). Therefore, the initially screened library was a good representation of the whole design space and already achieved the highest possible production.

### 3.2 DBTL Cycle 2: expansion of the original design space

ML analysis suggested that the optimal production possible considering the initial design space had already been found. In order to validate this prediction, top predicted designs by all the learning strategies were constructed. Figure 3B shows predicted and measured production of the top 7 designs in the CR ranking. As expected, production of these strains did not significantly improve with respect to the BMP strain. Similar results were obtained with the top strains from the CO, PO and PR rankings, with production remaining within the BMP mean *±* 20% (Sup Figure 8).

In order to improve pCA production, the original design space had to be expanded, and permutation feature importance and SHAP values were used to guide the new designs. Permutation feature importance identifies the factors with the biggest influence on model performance by evaluating the decrease in model accuracy when the values of a factor are shuffled. Factor 5, representing the expression strength of C4H, was identified as the most relevant factor, followed by factor 4 (PAL expression) (Figure 4A, Sup Figure 10). Considering that the predicted top producers had C4H under the strongest promoter tested, and never chose the weaker promoter for PAL, we hypothesized that higher expression of these genes could lead to higher production. This was confirmed by the SHAP values, a technique for explainable ML based on game theory, that not only identifies significant factors, but also determines how they affect the model output [27]. For all the training strategies used (except MLR model with the producer dataset), the highest positive impact on model output (*i*.*e*. production) was caused by expressing C4H and PAL under the strongest promoter tested. Similarly, the highest negative impact was caused by expression of C4H and PAL under weaker promoters (Figure 4B, Sup Figure 11). The importance of these genes was confirmed by substituting the promoters of PAL and/or C4H by weak promoters in the BMP strain and the best strain in the CR ranking (CR1). In both cases, strains with lower expression of PAL and/or C4H showed significantly reduced pCA production (Figure 4C). Besides, although the effect of different expression levels of CPR could not be assessed due to unsuccessful cassette construction, changing the promoter of CPR in the BMP and CR1 strains did not significantly change pCA production (Sup Figure 9).

**Figure 4:**
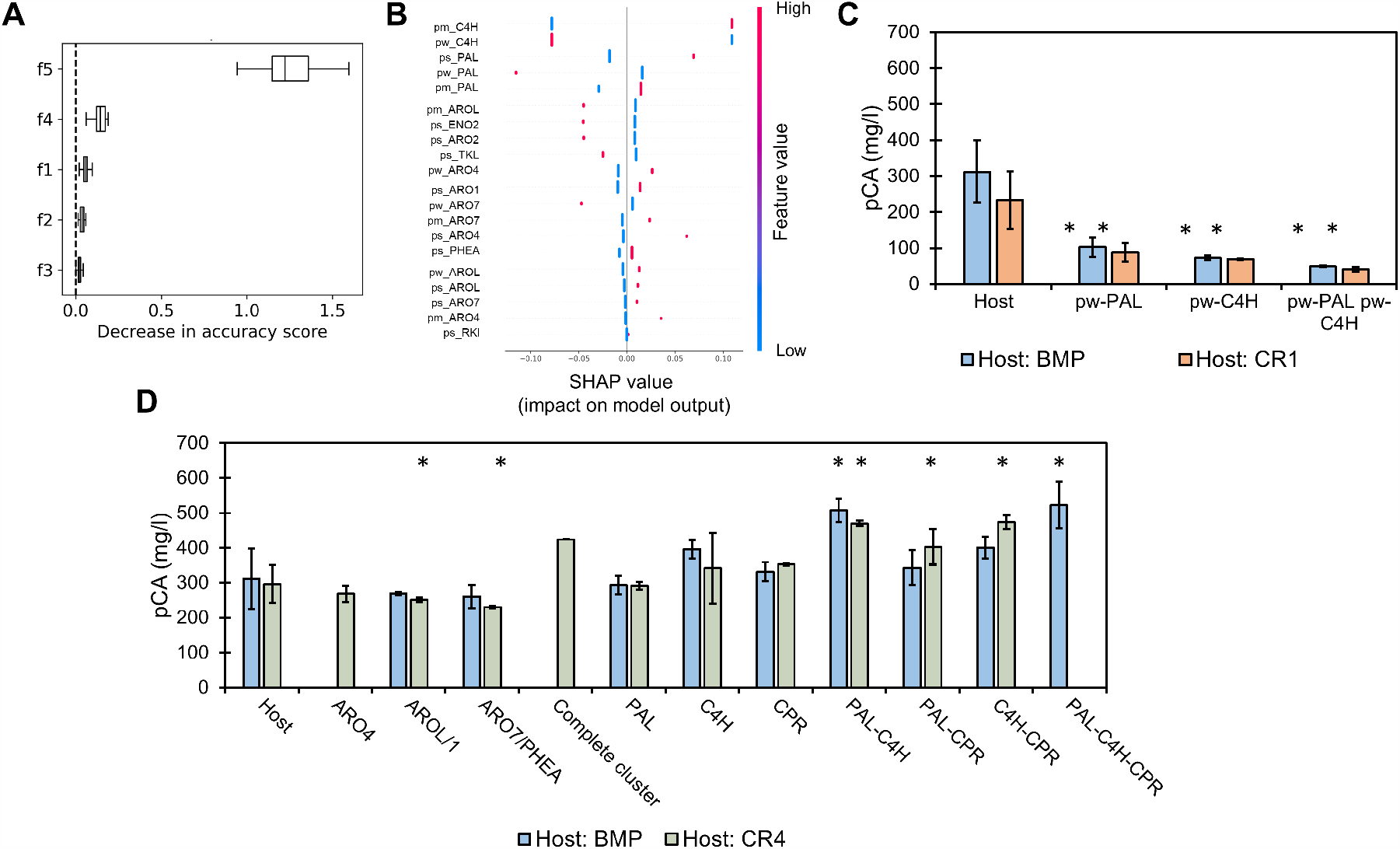
**A**. Representative example of feature selection results, where f1, f2, f3, f4 and f5 refer to factors 1 to 5. **B**. Representative example of SHAP values, where ps, pm and pw refer to strong, medium and weak promoters, respectively. **C**. Effect of substituting promoters of PAL and/or C4H by the weakest alternative (pw) in two different hosts: best measured producer (BMP) and predicted top producer by the *complete recurrent* strategy (CR1). **D**. Effect of integration of double copies of genes in two different hosts: BMP and the constructed predicted top producer by the *complete recurrent* strategy (CR4). Significant differences respect to each host are indicated by *. Colonies with double copies of ARO4 and the complete gene cluster were not obtained in the BMP host. Colonies with double copies of PAL-C4H-CPR were not obtained in the CR4 host and a single colony was obtained with the correct integration of the complete gene cluster.

To further increase the expression of the genes, strains with double copies of each of the genes were created using BMP and the best constructed strain from the CR ranking (CR4) as hosts. Positive colonies containing double copies of ARO4 and ARO4-AROL-ARO7-PAL-C4H-CPR in the BMP host and PAL-C4H-CPR in the CR4 host were not found. As expected, when extra copies of factors 1 (ARO4), 2 (AROL or ARO1) and 3 (ARO7 or PHEA) were integrated, production of pCA did not significantly change (Figure 4D). However, production did not significantly increase either when double copies of PAL or C4H were integrated. Even though average production increased with a double copy of C4H, this change was not significant (Figure 4D). The integration of a double copy of the complete gene cluster was only achieved in one colony of the CR4 host, and its production was similar to strains with an extra copy of C4H. In both hosts, double copies of PAL and C4H resulted in significantly increased production (63% in BMP and 58% in CR4). Besides, significantly increased production was also found when double copies of PAL-CPR (36%) and C4H-CPR (60%) were expressed in CR4; and PAL-C4H-CPR were expressed in BMP (68%) (Figure 4D).

The observed increase in pCA production, only obtained when expanding the original design space, confirmed that the original space had been sufficiently sampled and validated feature importance as a strategy to guide its expansion.

## 4 Discussion

Accelerating the design of industrially relevant strains is crucial to transition to a bio-based economy. In order to exploit the full potential of microorganisms, combinatorial optimization of metabolic pathways is required. However, this involves the construction and testing of an exponentially growing number of strains which becomes unfeasible [31]. Alternatively, the solution space can be sampled following a rational or randomized approach. Statistical design of experiments reduces the number of strains to build and test while maximizing the information gained about the complete solution space. However, it requires the construction of specific strains and it is sensitive to experimental limitations: information is lost when a strain cannot be built [32]. As shown here, ML presents an alternative to learn from randomly generated libraries of strains which is robust to missing data. Besides, when ML is used, libraries can be flexibly designed to include factors with different number of levels based on prior knowledge. We used factor 1 to explore genes that could influence pCA production assigning 7 levels to this factor. Instead, we assigned 3 levels to factors 4, 5 and 6, aiming to fine-tune the expression of the required heterologous genes. We used 4 levels for factors 2 and 3 to simultaneously test the effect of homologous ORFs from different origins and tune the expression of one of them. The robustness and flexibility of the ML approach was also shown when some of the designed levels could not be implemented experimentally. Although the library design space was reduced from 3024 members to 672, the relationship between the remaining levels could still be efficiently explored.

Another challenge to combinatorial pathway optimization is the need of characterization of genetic parts that ensures that the solution space is sufficiently explored. This is especially important when we aim at fine-tuning the expression levels of pathway genes [5, 8, 10, 12]. Although effort is taken to appropriately characterize how regulatory elements affect gene expression, this is seldom achieved as the effect of the regulatory elements is ORF-dependent [33]. Alternatively, regulatory elements can be treated as categorical variables reducing the impact of the characterization data [8]. This approach allowed us to include non-characterized promoters as members of the library and avoid a further decrease of the design space size. Besides, the use of categorical variables does not limit factor’s levels to differences in expression strength. As shown here, factors might include levels that represent differences in expression but also different ORFs, broadening the scope of ML-guided pathway optimization to the selection of genes from different origins or alternative over-expression targets.

A limitation to the use of ML is the requirement for sufficient and quality data for training [4]. We showed that including non-producers as part of the training set is not required to find top producing strains but improves predictions of poor producers which helps ensuring that the design space has been sufficiently sampled. This is especially important when the top producer is already present in the training data. Frequently, ML algorithms are trained with data representing circa 5% of the library design space. Although we trained the ML algorithms with data representing 13.5% of the library, the effect of reducing the amount of training data was tested. When 40% of the available training data was used for training with stratification (equivalent to 5.4% of the library space), coefficients of determination remained above 0.6 for all the models but MLR regardless of the dataset used (Sup Figure 12, Table 2). When the amount of data used for training decreased, stratification during training improved the mean R^2^ and reduced its standard deviation. Stratification allowed the classification of samples based on production. Therefore, a sufficient number of samples from each category should be present in the training data. As shown here, this can be achieved using a screening before sequencing approach, which allowed an efficient exploration of the design space and reduced the chance of sequencing duplicate designs.

**Table 2:**
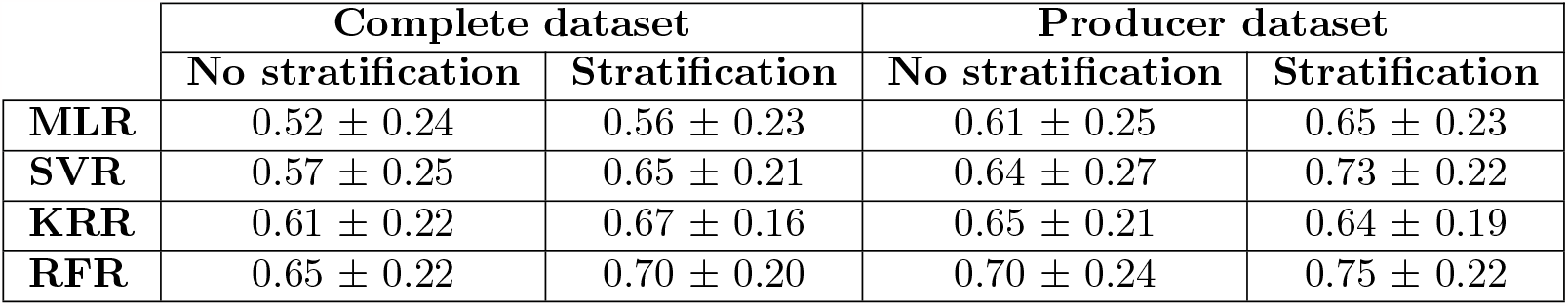
R^2^ with 5.4% of the library as training data size.

|  | Complete dataset |  | Producer dataset |  |
| --- | --- | --- | --- | --- |
|  | No stratification | Stratification | No stratification | Stratification |
| **MLR** | $0.52 \pm 0.24$ | $0.56 \pm 0.23$ | $0.61 \pm 0.25$ | $0.65 \pm 0.23$ |
| **SVR** | $0.57 \pm 0.25$ | $0.65 \pm 0.21$ | $0.64 \pm 0.27$ | $0.73 \pm 0.22$ |
| **KRR** | $0.61 \pm 0.22$ | $0.67 \pm 0.16$ | $0.65 \pm 0.21$ | $0.64 \pm 0.19$ |
| **RFR** | $0.65 \pm 0.22$ | $0.70 \pm 0.20$ | $0.70 \pm 0.24$ | $0.75 \pm 0.22$ |

ML algorithms cannot extrapolate, they cannot predict the performance of strains with factor levels different from those used during training [7]. Still, they can be used to determine whether the best producer from the library is already present in the training data and justify the expansion of the original design space. When this is required, we proposed the use of feature importance and SHAP values to guide this expansion and point at the most relevant factors which, in this case, lead to a 68% improvement in pCA production. Notably, while feature importance only points as the significant factors, SHAP values provide additional information regarding how the factor’s levels influence the model output[27].

The highest titers of pCA measured in this study were 0.51 *±* 0.03 and 0.52 *±* 0.06 g/l obtained using the BMP strain with additional copies of PAL-C4H or PAL-C4H-CPR. These strains were cultivated in 96DWP with 20 g/l of glucose resulting on 0.03 g/g pCA yield on glucose. However, higher titers and yields of pCA have been reported. Rodriguez et al. obtained 1.96 g/l of pCA (0.04 g/g) by expressing AROL, feedback resistant variants of ARO4 and ARO7, eliminating competing metabolic pathways and using synthetic fed-batch media [18]. Production was further improved by Liu et al. combining the TAL and PAL pathways and including a phosphoketolase pathway to increase E4P availability. This strain produced 3.1 g/l in shake flasks and up to 12.5 g/l in bioreactors operated as fed-batch with a maximum yield of 0.15 g/g [14]. Considering this results, production of our developed strains could be further improved in next cycles that focus on gene deletions and bio-process optimization.

This study is an example of how ML-guided DBTL cycles can accelerate the generation of efficient strains. We showed the robustness of this approach to experimental limitations and its flexibility regarding design, which can be expanded beyond the traditional tuning of gene expression. We propose a screening before sequencing approach to allow for stratification during training, especially important for small datasets. Furthermore, we showed how feature importance and SHAP values can be used to expand the original design space and further improve strain performance.

## 5 Acknowledgment

This project was founded by the Netherlands Organization for Scientific Research (NWO; project number GSGT.2019.008) and the European Union’s Horizon 2020 research and innovation program under grant agreement 814408 (Shikifactory100) . Additionally we would like to thank Moniek Jonkers for her help with the laboratory automation workflows and Lieke Meijvogel, Sharina Chander, Judith Vis, Sylvana Suisse, Wibo B. van Scheppingen and Leon Coulier for execution and support with the analytical workflows.

## References

[1] Kim, G. B. et al. “Metabolic engineering for sustainability and health”. In: Trends in Biotechnology 41.3 (2023), 425–451. doi: 10.1016/J.TIBTECH.2022.12.014.

[2] Hodgman, C. E. and Jewett, M. C. “Cell-free synthetic biology: Thinking outside the cell”. In: Metabolic Engineering 14.3 (2012), 261–269. doi: 10.1016/J.YMBEN.2011.09.002.

[3] Carbonell, P. et al. “An automated Design-Build-Test-Learn pipeline for enhanced microbial production of fine chemicals”. In: Communications Biology 1.1 (2018), 66. doi: 10.1038/s42003.018.0076.9.

[4] Lawson, C. et al. “Machine learning for metabolic engineering: A review”. In: Metabolic Engineering (2020). doi: 10.1016/j.ymben.2020.10.005.

[5] HamediRad, M. et al. “Towards a fully automated algorithm driven platform for biosystems design”. In: Nature Communications 10.1 (2019), 1–10. doi: 10.1038/s41467.019.13189.z.

[6] Liao, X. et al. “Artificial intelligence: a solution to involution of design–build–test–learn cycle”. In: Current Opinion in Biotechnology 75 (2022), 102712. doi: 10.1016/J.COPBIO.2022.102712.

[7] Oyetunde, T. et al. “Leveraging knowledge engineering and machine learning for microbial biomanufacturing”. In: Biotechnology Advances 36.4 (2018), 1308–1315. doi: 10.1016/j.biotechadv.2018.04.008.

[8] Zhang, J. et al. “Combining mechanistic and machine learning models for predictive engineering and optimization of tryptophan metabolism”. In: Nature Communications 11.1 (2020), 1–13. doi: 10.1038/s41467.020.17910.1.

[9] Jervis, A. J. et al. “Machine Learning of Designed Translational Control Allows Predictive Pathway Optimization in Escherichia coli”. In: ACS Synthetic Biology 8.1 (2019), 127–136. doi: 10.1021/acssynbio.8b00398.

[10] Zhou, Y. et al. “MiYA, an efficient machine-learning workflow in conjunction with the YeastFab assembly strategy for combinatorial optimization of heterologous metabolic pathways in Saccharomyces cerevisiae”. In: Metabolic Engineering 47 (2018), 294–302. doi: 10.1016/J.YMBEN.2018.03.020.

[11] Keun Kang, C. et al. “Machine learning-guided prediction of potential engineering targets for microbial production of lycopene”. In: Bioresource Technology (2022), 128455. doi: 10.1016/J.BIORTECH.2022.128455.

[12] Opgenorth, P. et al. “Lessons from Two DesignBuildTestLearn Cycles of Dodecanol Production in Escherichia coli Aided by Machine Learning”. In: ACS Synth. Biol (2019). doi: 10.1021/acssynbio.9b00020.

[13] Radivojević, T. et al. “A machine learning Automated Recommendation Tool for synthetic biology”. In: Nature Communications 11.1 (2020), 1–14. doi: 10.1038/s41467.020.18008.4.

[14] Liu, Q. et al. “Rewiring carbon metabolism in yeast for high level production of aromatic chemicals”. In: Nature Communications 10.1 (2019), 1–13. doi: 10.1038/s41467.019.12961.5.

[15] Kanehisa, M. and Goto, S. “KEGG: Kyoto Encyclopedia of Genes and Genomes”. In: Nucleic Acids Research 28.1 (2000), 27–30. doi: 10.1093/NAR/28.1.27.

[16] Braus, G. H. “Aromatic amino acid biosynthesis in the yeast Saccharomyces cerevisiae: A model system for the regulation of a eukaryotic biosynthetic pathway”. In: Microbiological Reviews 55.3 (1991), 349–370. doi: 10.1128/mmbr.55.3.349-370.1991.

[17] Koopman, F. et al. “De novo production of the flavonoid naringenin in engineered Saccharomyces cerevisiae”. In: Microbial Cell Factories 11.1 (2012), 1–15. doi: 10.1186/1475.2859.11.155.

[18] Rodriguez, A. et al. “Establishment of a yeast platform strain for production of p-coumaric acid through metabolic engineering of aromatic amino acid biosynthesis”. In: Metabolic Engineering 31 (2015), 181?188–188. doi: 10.1016/j.ymben.2015.08.003.

[19] Jendresen, C. B. et al. “Highly active and specific tyrosine ammonia-lyases from diverse origins enable enhanced production of aromatic compounds in bacteria and Saccharomyces cerevisiae”. In: Applied and Environmental Microbiology 81.13 (2015), 4458–4476. doi: 10.1128/AEM.00405.15.

[20] Young, E. M., Gordon, D. B., and Voigt, C. Composability and design of parts for large-scale pathway engineering in yeast. 2015.

[21] Roubos, J. A. and VAN Noel, N. M. E. A method for achieving improved polypeptide expression. 2007.

[22] Verwaal, R. et al. “CRISPR/Cpf1 enables fast and simple genome editing of Saccharomyces cerevisiae”. In: Yeast 35.2 (2018), 201–211. doi: 10.1002/YEA.3278. url: https://onlinelibrary.wiley.com/doi/full/10.1002/yea.3278%20 https://onlinelibrary.wiley.com/doi/abs/10.1002/yea.3278%20 https://onlinelibrary.wiley.com/doi/10.1002/yea.3278.

[23] Ciurkot, K. et al. “Efficient multiplexed gene regulation in Saccharomyces cerevisiae using dCas12a”. In: Nucleic Acids Research 49.13 (2021), 7775–7790. doi: 10.1093/NAR/GKAB529. url: https://dx.doi.org/10.1093/nar/gkab529.

[24] Gietz, R. D. et al. “Studies on the transformation of intact yeast cells by the LiAc/SS-DNA/PEG procedure”. In: Yeast 11.4 (1995), 355–360. doi: 10.1002/YEA.320110408. url: https://onlinelibrary.wiley.com/doi/full/10.1002/yea.320110408%20https://onlinelibrary.wiley.com/doi/abs/10.1002/yea.320110408%20https://onlinelibrary.wiley.com/doi/10.1002/yea.320110408.

[25] Dekker, W. J. et al. “Anaerobic growth of Saccharomyces cerevisiae CEN.PK113-7D does not depend on synthesis or supplementation of unsaturated fatty acids”. In: FEMS Yeast Research 19.6 (2019), 60. doi: 10.1093/FEMSYR/FOZ060. url: https://dx.doi.org/10.1093/femsyr/foz060.

[26] Wouters, B. et al. “Automated Segmented-Flow Analysis - NMR with a Novel Fluoropolymer Flow Cell for High-Throughput Screening”. In: Analytical Chemistry 94.44 (2022), 15350–15358. doi: 10.1021/ACS.ANALCHEM.2C03038/ASSET/IMAGES/LARGE/AC2C030380005.JPEG. url: https://pubs.acs.org/doi/full/10.1021/acs.analchem.2c03038.

[27] Lundberg, S. M., Allen, P. G., and Lee, S.-I. “A Unified Approach to Interpreting Model Predictions”. In: Advances in Neural Information Processing Systems 30 (2017). url: https://github.com/slundberg/shap.

[28] Pedregosa, F. et al. “Scikit-learn: Machine Learning in Python”. In: Journal of Machine Learning Research 12 (2011), 2825–2830.

[29] Gething, M. J. and Davidson, B. E. “Chorismate mutase/prephenate dehydratase from Escherichia coli K12. Effects of chemical modification on the enzymic activities and allosteric inhibition”. In: European journal of biochemistry 78.1 (1977), 111–117. doi: 10.1111/J.1432-1033.1977.TB11719.X.

[30] Sampathkumar, P. and Morrison, J. F. “Chorismate mutase-prephenate dehydrogenase from Escherichia coli. Purification and properties of the bifunctional enzyme”. In: Biochimica et biophysica acta 702.2 (1982), 204–211. doi: 10.1016/0167-4838(82)90504-0.

[31] Brown, S. R. et al. “Design of Experiments Methodology to Build a Multifactorial Statistical Model Describing the Metabolic Interactions of Alcohol Dehydrogenase Isozymes in the Ethanol Biosynthetic Pathway of the Yeast Saccharomyces cerevisiae”. In: ACS Synthetic Biology 7.7 (2018), 1676–1684. doi: 10.1021/acssynbio.8b00112. url: http://pubs.acs.org/doi/10.1021/acssynbio.8b00112.

[32] Young, E. M. et al. “Iterative algorithm-guided design of massive strain libraries, applied to itaconic acid production in yeast”. In: Metabolic Engineering 48 (2018), 33–43. doi: 10.1016/j.ymben.2018.05.002. url: https://linkinghub.elsevier.com/retrieve/pii/S1096717618301186.

[33] Carquet, M., Pompon, D., and Truan, G. “Transcription interference and ORF nature strongly affect promoter strength in a reconstituted metabolic pathway”. In: Frontiers in Bioengineering and Biotechnology 3.FEB (2015), 132035. doi: 10.3389/FBIOE.2015.00021/ABSTRACT.

