## Supplementary material for "Machine learning-guided optimization of p-coumaric acid production in yeast": Sup. Figure

**TABLE OF CONTENTS**

| **Supplementary Tables** |  |
| --- | --- |
| Supplementary Table 1: Promoters, ORF and terminator sequences | 2 |
| Supplementary Table 2: Integration sites used for strains construction | 2 |
| Supplementary Table 3: List of strains used in this study | 2 |
| Supplementary Table 4: List of plasmids used in this study | 2 |
| Supplementary Table 5: List of primers used in this study | 2 |
| Supplementary Table 6: Features of sgRNA and crRNA design | 3 |
| **Supplementary Figures** |  |
| Supplementary Figure 1. Library design and construction | 4 |
| Supplementary Figure 2. Promoter-terminator characterization | 4 |
| Supplementary Figure 3. pCA production correct strain PAL library | 5 |
| Supplementary Figure 4. Model selection and training strategies | 5 |
| Supplementary Figure 5. Ranking of BMP, top 5 BMP and worst producers | 6 |
| Supplementary Figure 6. Genotype top 10 predicted strains all learning strategies | 7 |
| Supplementary Figure 7. Phenotype top 10 predicted strains all learning strategies | 8 |
| Supplementary Figure 8. Validation ML predictions | 9 |
| Supplementary Figure 9. Additional promoters CPR | 10 |
| Supplementary Figure 10. Feature importance | 10 |
| Supplementary Figure11. Effect of training data size on model accuracy | 11 |
| **Supplementary References** | 11 |

Supplementary Table 1: Promoters, ORFs and terminator sequences

See attached Excel file.

Supplementary Table 2: Integration sites used for strains construction.

| **Integration site** | **Chromosome** | **Location** | **Experiment** |
| --- | --- | --- | --- |
| INT1 | 15 | Non-coding region between NTR1 (YOR071c) and GYP1 (YOR070c) | Cas9 integration |
| INT69B |  | (Genbank: AEHG01000256.1) | DBTL Cycle 1 |
| INT72B |  | (Genbank: AEHG01000557.1) | DBTL Cycle 2 |

Supplementary Table 3: List of strains used in this study.

See attached Excel file.

Supplementary Table 4: List of plasmids used in this study.

| Plasmid | Genotype | Reference | Addgene |
| --- | --- | --- | --- |
| pCSN061 | *CEN6/ARS TRP1* KanMX bla Kl11p-SpCas9-GND2t | (2) | # 101725 |
| pRN1120 | 2µm NatMX amp^R^ | (2) | # 101750 |

Supplementary Table 5: List of primers used in this study.

| **Name** | **Sequence** | **Purpose** |
| --- | --- | --- |
| pSH001 | CTTATCGATACCGTCGACCTCGAGGGGGGGCCCGGTACCCAGCTTTTGTTCCGCGGTCTTTGAAAAGATAATG | Amplification of Cas9 sgRNA expression cassette (RV) |
| pSH002 | AAAATACAACAAATAAAAAACACTCAATGACCTGACCATTTGATGGAGTTCCGCGGAGACATAAAAAAC | Amplification of Cas9 sgRNA expression cassette (FW) |
| pSH003 | AACTCCATCAAATGGTCAGG | Amplification of Cas9 sgRNA recipient plasmid pRN1120 (FW) |
| pSH004 | AACAAAAGCTGGGTACCGGG | Amplification of Cas9 sgRNA recipient plasmid pRN1120 (RV) |
|  |  | Amplification of INT69B left flank (RV) with homology to conX |
|  |  | Amplification of INT69B right flank (FW) with homology to conX |
| Add primers used to amplify cassettes |  |  |

Supplementary Table 5: sgRNA design.

| **Target Locus** | **GBlock sequence** |
| --- | --- |
| INT69 | TACCCAGCTTTTGTTCCGCGGTCTTTGAAAAGATAATGTATGATTATGCTTTCACTCATATTTATACAGAAACTTGATGTTTTCTTTCGAGTATATACAAGGTGATTACATGTACGTTTGAAGTACAACTCTAGATTTTGTAGTGCCCTCTTGGGCTAGCGGTAAAGGTGCGCATTTTTTCACACCCTACAATGTTCTGTTCAAAAGATTTTGGTCAAACGCTGTAGAAGTGAAAGTTGGTGCGCATGTTTCGGCGTTCGAAACTTCTCCGCAGTGAAAGATAAATGATCAAACAACAGTTCCGTAACCGGTTTTAGAGCTAGAAATAGCAAGTTAAAATAAGGCTAGTCCGTTATCAACTTGAAAAAGTGGCACCGAGTCGGTGGTGCTTTTTTTGTTTTTTATGTCTCCGCGGAACTCCATCAAATGG |
| INT72 | TACCCAGCTTTTGTTCCGCGGTCTTTGAAAAGATAATGTATGATTATGCTTTCACTCATATTTATACAGAAACTTGATGTTTTCTTTCGAGTATATACAAGGTGATTACATGTACGTTTGAAGTACAACTCTAGATTTTGTAGTGCCCTCTTGGGCTAGCGGTAAAGGTGCGCATTTTTTCACACCCTACAATGTTCTGTTCAAAAGATTTTGGTCAAACGCTGTAGAAGTGAAAGTTGGTGCGCATGTTTCGGCGTTCGAAACTTCTCCGCAGTGAAAGATAAATGATCGATGCTGACACAATTAATGTGTTTTAGAGCTAGAAATAGCAAGTTAAAATAAGGCTAGTCCGTTATCAACTTGAAAAAGTGGCACCGAGTCGGTGGTGCTTTTTTTGTTTTTTATGTCTCCGCGGAACTCCATCAAATGG |

Primer homology

SNR52p and ATG

Target sequence

Sup4T

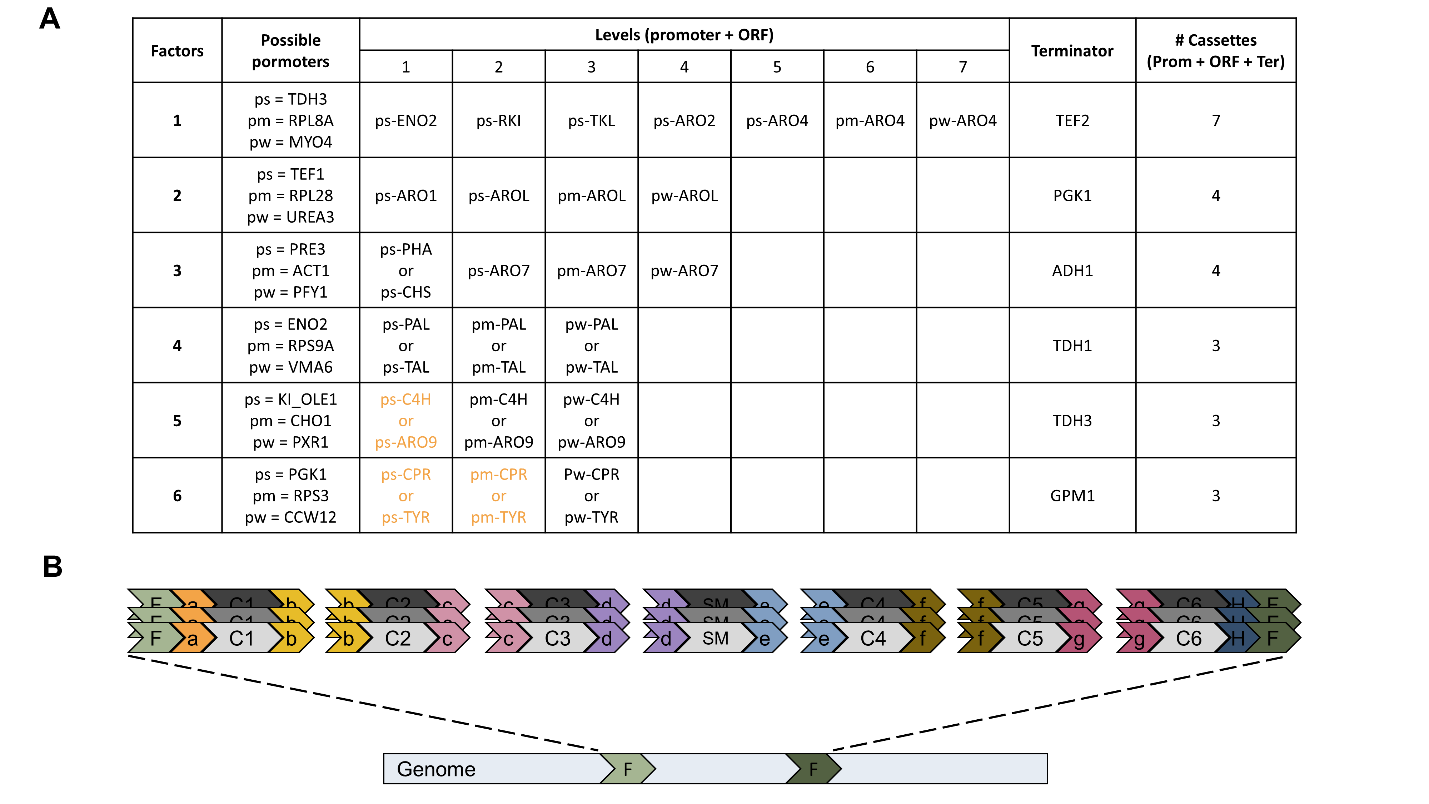

**Figure Sup. 1:** **A**. Cassettes used for library transformation. A cassette is a combination of promoter, ORF and terminator. Cassettes containing promoter-ORF combinations shown in orange could not be obtained. **B**. Schematic representation of the integration of a gene cluster. Connector sequences a to g represent homology regions for *in vivo* recombination of cassettes (C1 to C6) and the selection marker cassette (SM); flank sequences homologous to the genome integration site are shown as F.

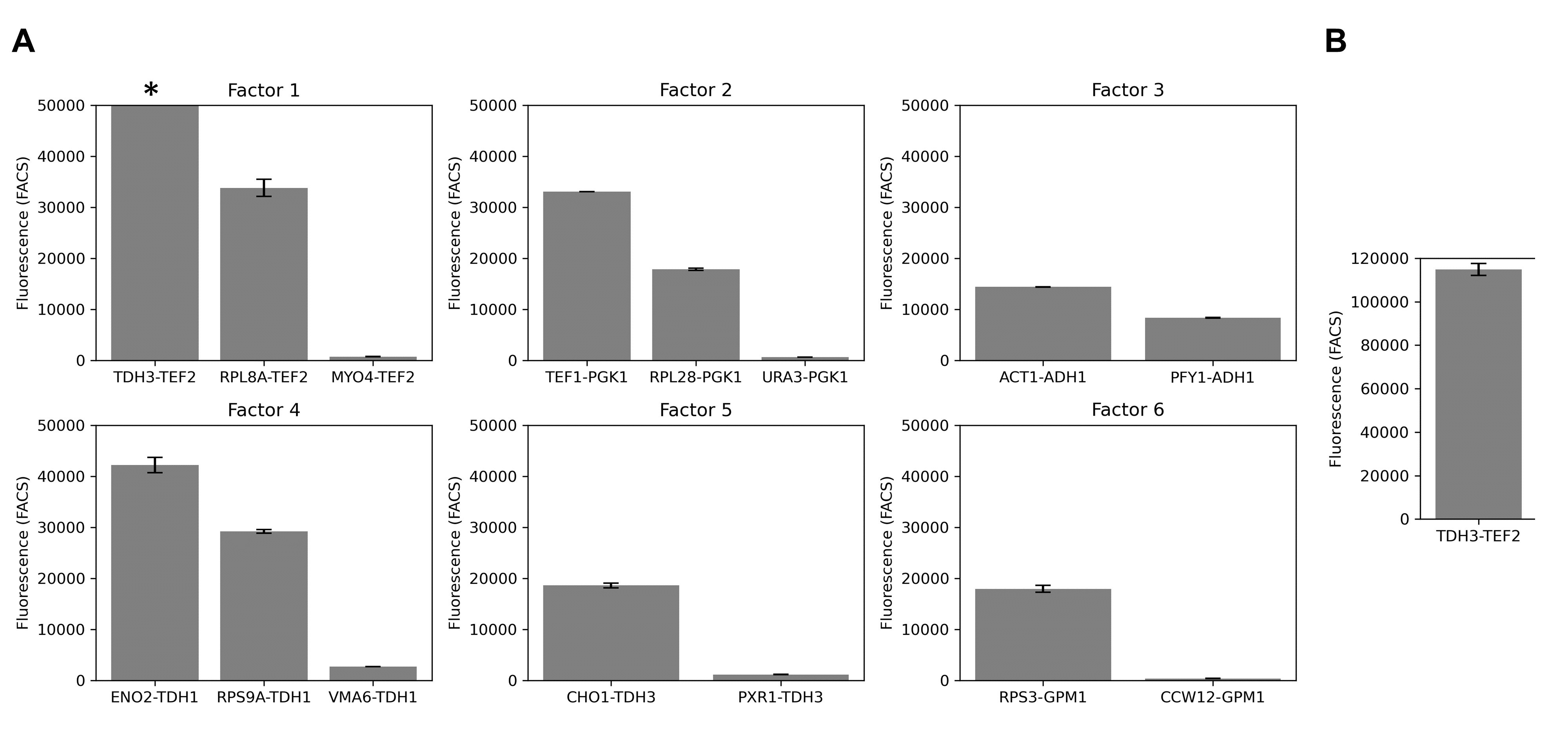
**Figure Sup. 2:** Promoter-terminator characterization by GFP fluorescence measured using fluorescence activated cell sorting (FACS) (**A**). Zoom-out for the fluorescence values (**B**).

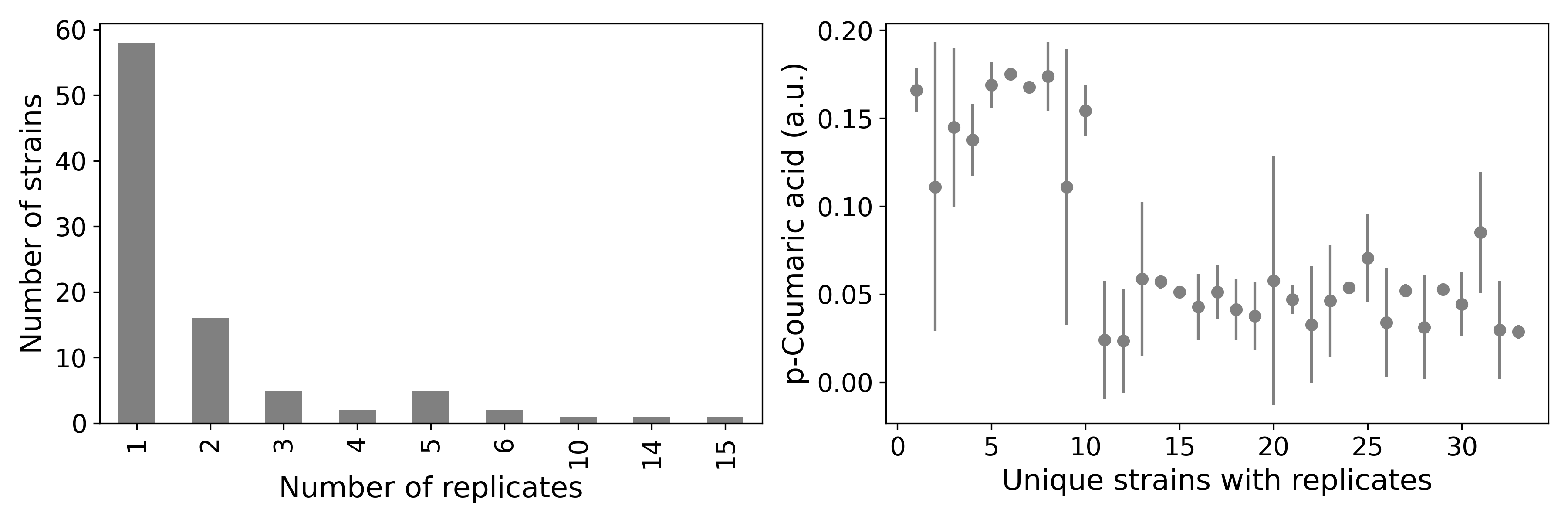
**Figure Sup. 3:** Characterization of the correct sequences from the PAL library. Frequency of strains with the same designs (left) and average _ρ_CA production of designs with replicates (right).

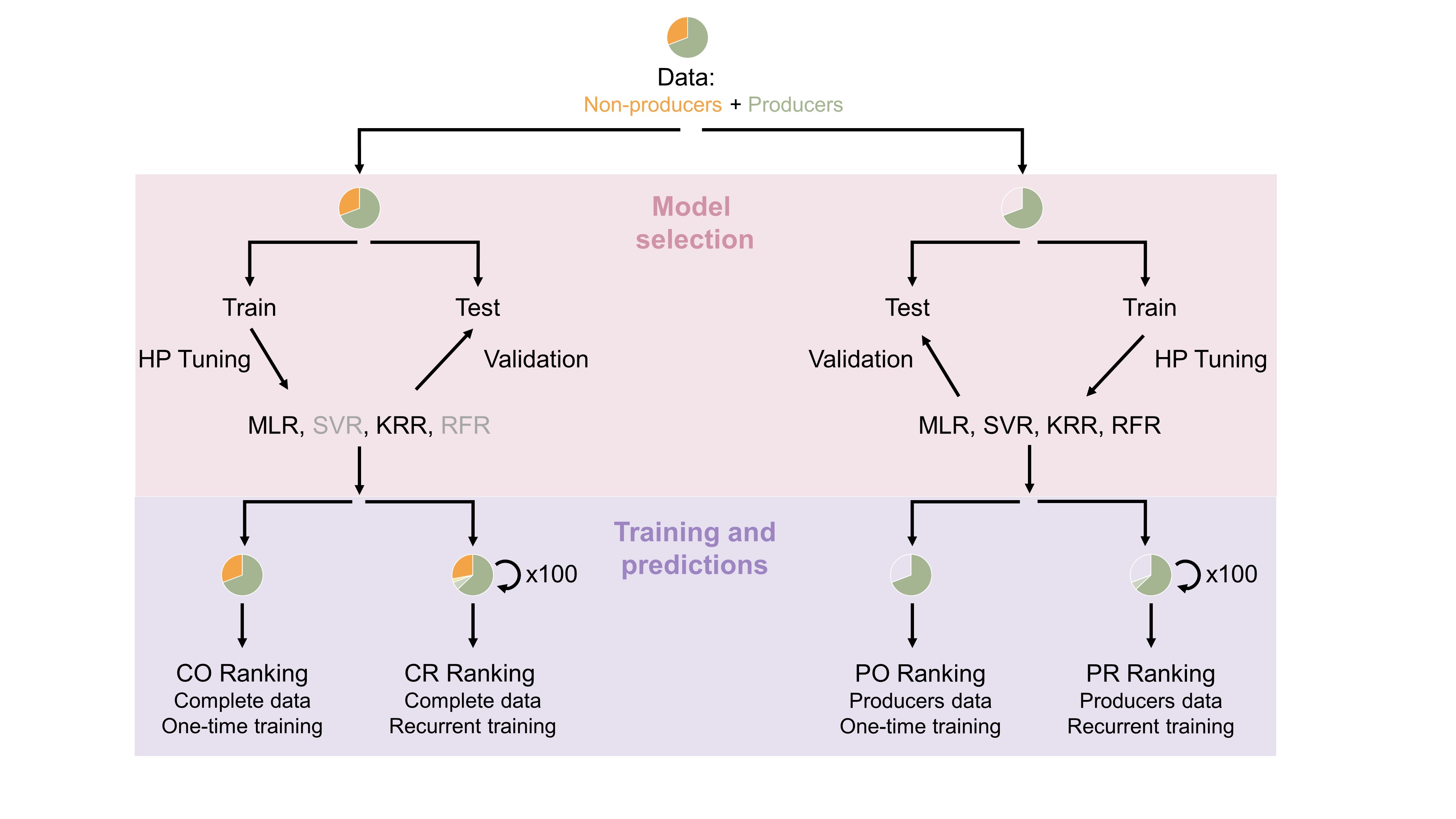

**Figure Sup. 4:** Model selection and training strategies. Genotype and production data were divided in two datasets: the *complete* and *producer* datasets that differ in the inclusion of data from nonproducers. Each dataset was used for hyper-parameter (HP) tuning of four ML models: multiple linear regressor (MLR), support vector regressor (SVR), kernel ridge regressor (KRR) and random forest regressor (RFR). Accuracy of models with optimal HP was evaluated on the test sets and is shown in the table. For each dataset, two learning strategies were applied: *one train training*, where all the data was used for training and *recurrent training*, where 90% of the training data was iteratively used for training.

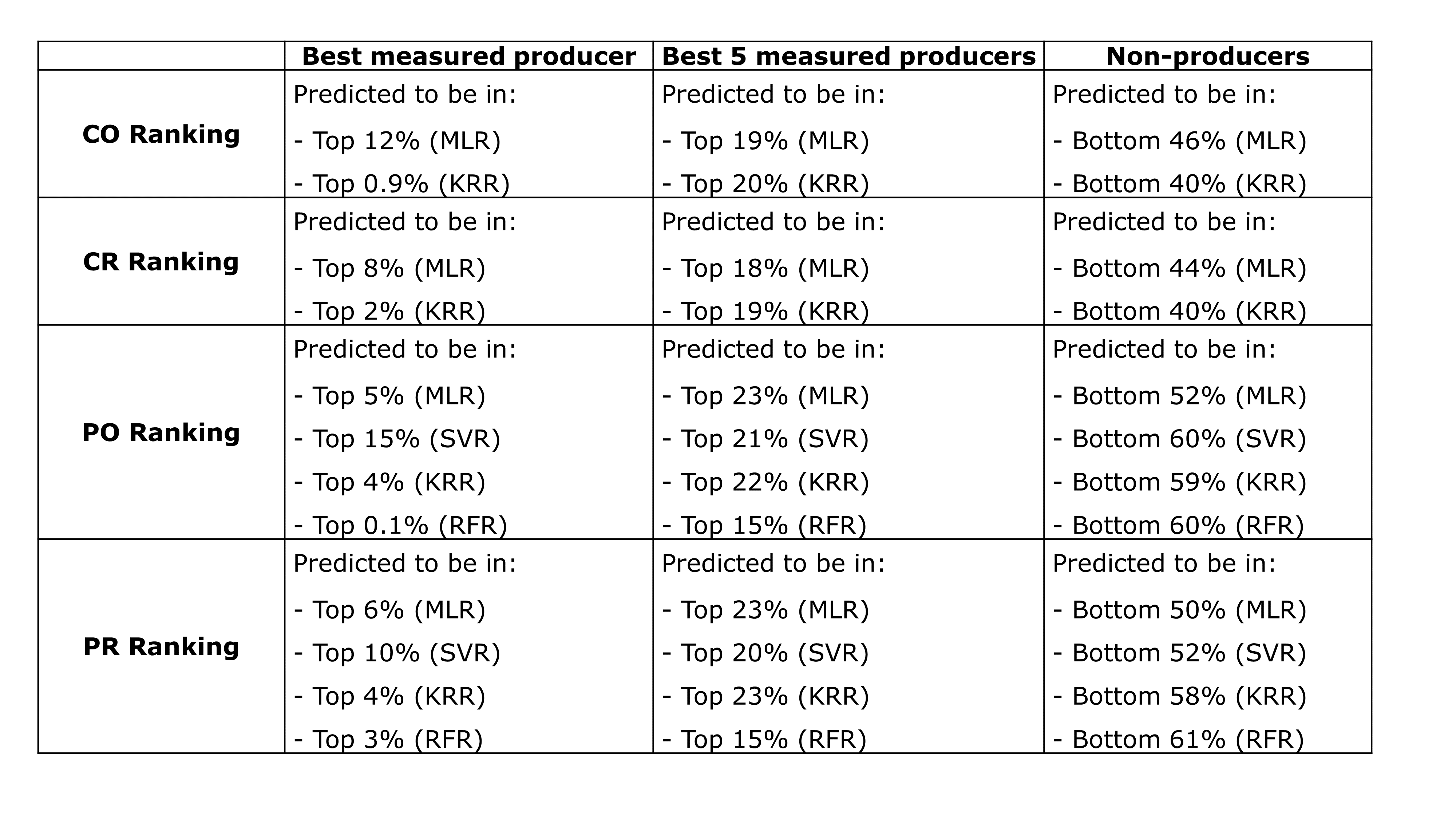

**Figure Sup. 5:** Ranking of top measured producer, top 5 measured producers and non-producers based on four different training strategies. Results are given per model used in each strategy: MLR, multiple linear regressor; KRR, kernel ridge regressor; SVR, support vector regressor; RFR, random forest regressor. *Ranking of non-produces excludes design 560 which is predicted to produce by all models independently of the training strategy. The BMP was ranked in the top 0.1% to 15% depending on the model used and regardless of the training strategy. Similarly, the 5-BMPs were always predicted to be, at least, in the top 22% of the library. Measured non-producers were ranked in the bottom 46% or 60% of the library depending on the dataset used for training (*complete* or *producers* respectively). Therefore, including non-producers during training did not change predictions of top producers, but improved predictions of non-producers, ensuring correct coverage of the complete library by the ML predictions.

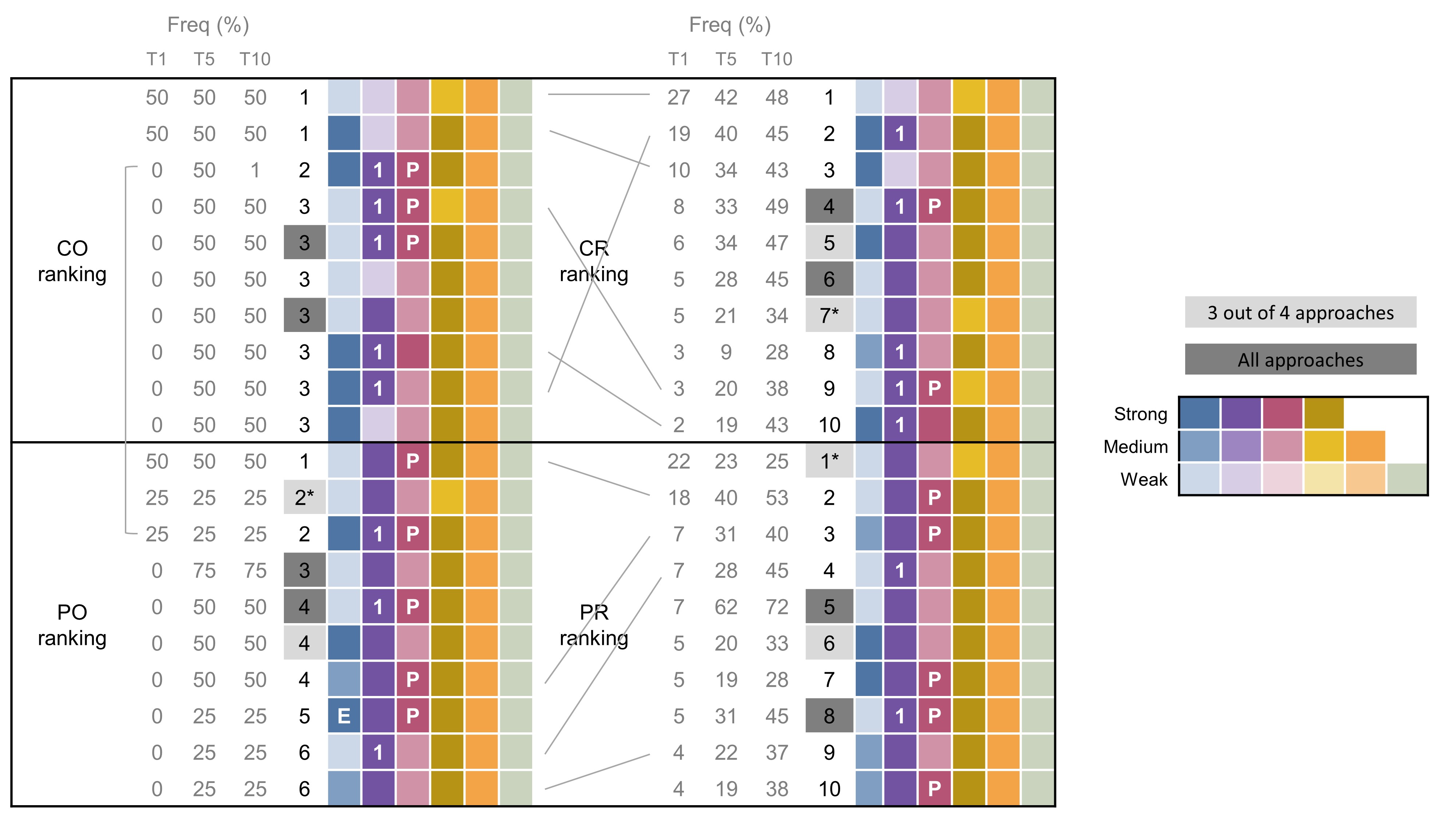

**Figure Sup. 6:** Summary of top 10 predicted producers by each learning strategy. Designs are ranked based on the frequency (Freq.) they are chosen as top 1 (T1), top 5 (T5) or top 10 (T10) by different models or data points included during training. Factor 1 refers to ARO4 except an E is shown (ENO1), factor 2 refers to AROL except a 1 is shown (ARO1), factor 3 refers to ARO7 unless a P is shown (PHEA), factor 4 refers to PAL, factor 5 refers to C4H and factor 6 refers to CPR. Predicted designs shared by different learning strategies are linked by lines or highlighted in grey. * indicates designs equal to the best measured producer strain (BMP strain).

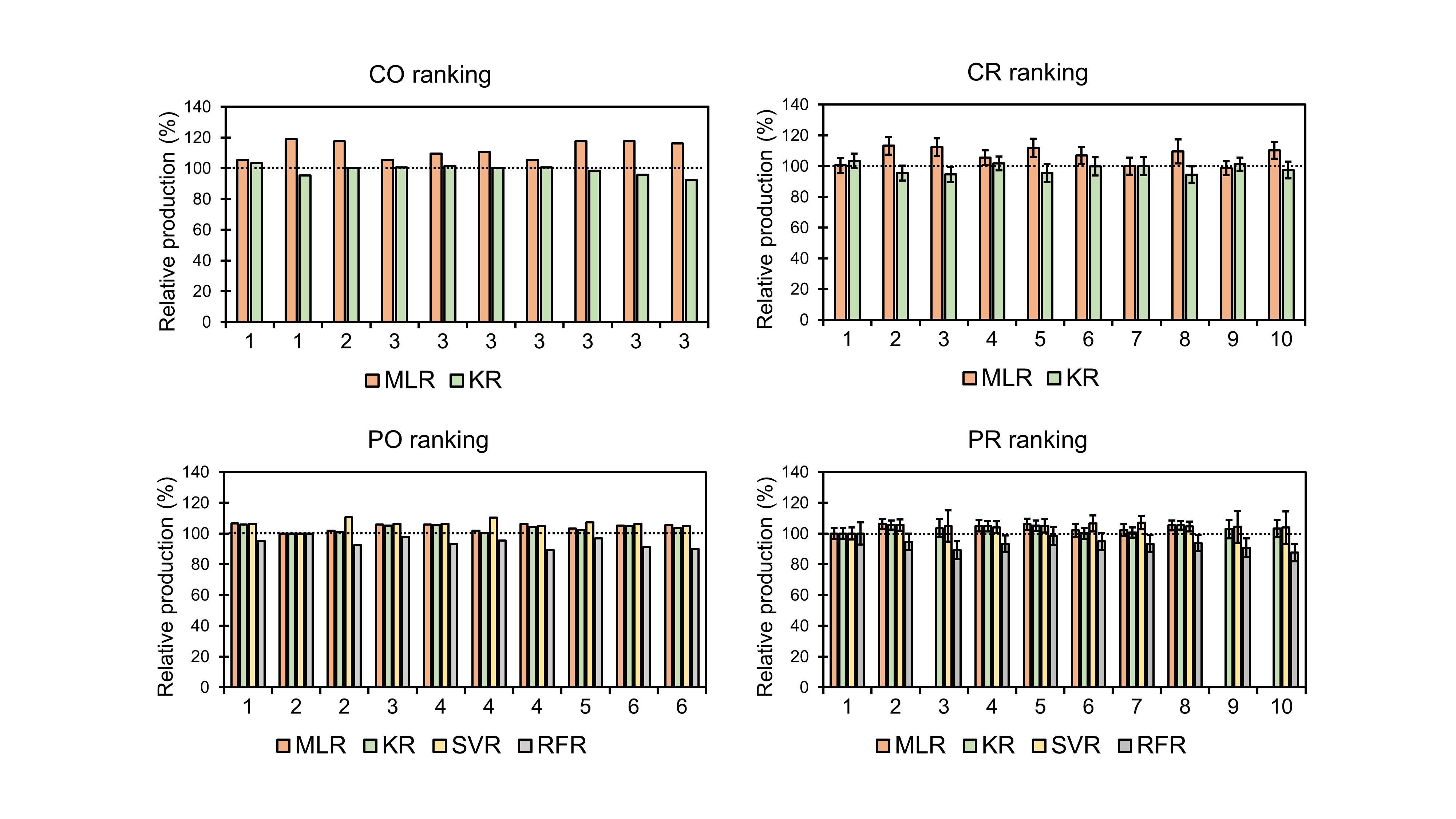

**Figure Sup. 7:** Predicted _ρ_CA production by the top 10 ranked strains found using four different leaning strategies. Production relative to the predicted production of the top measured producer is shown. CO, complete dataset with one time training; CR, complete dataset with recurrent training; PR, producers dataset with one time training; PR producers dataset with recurrent training.

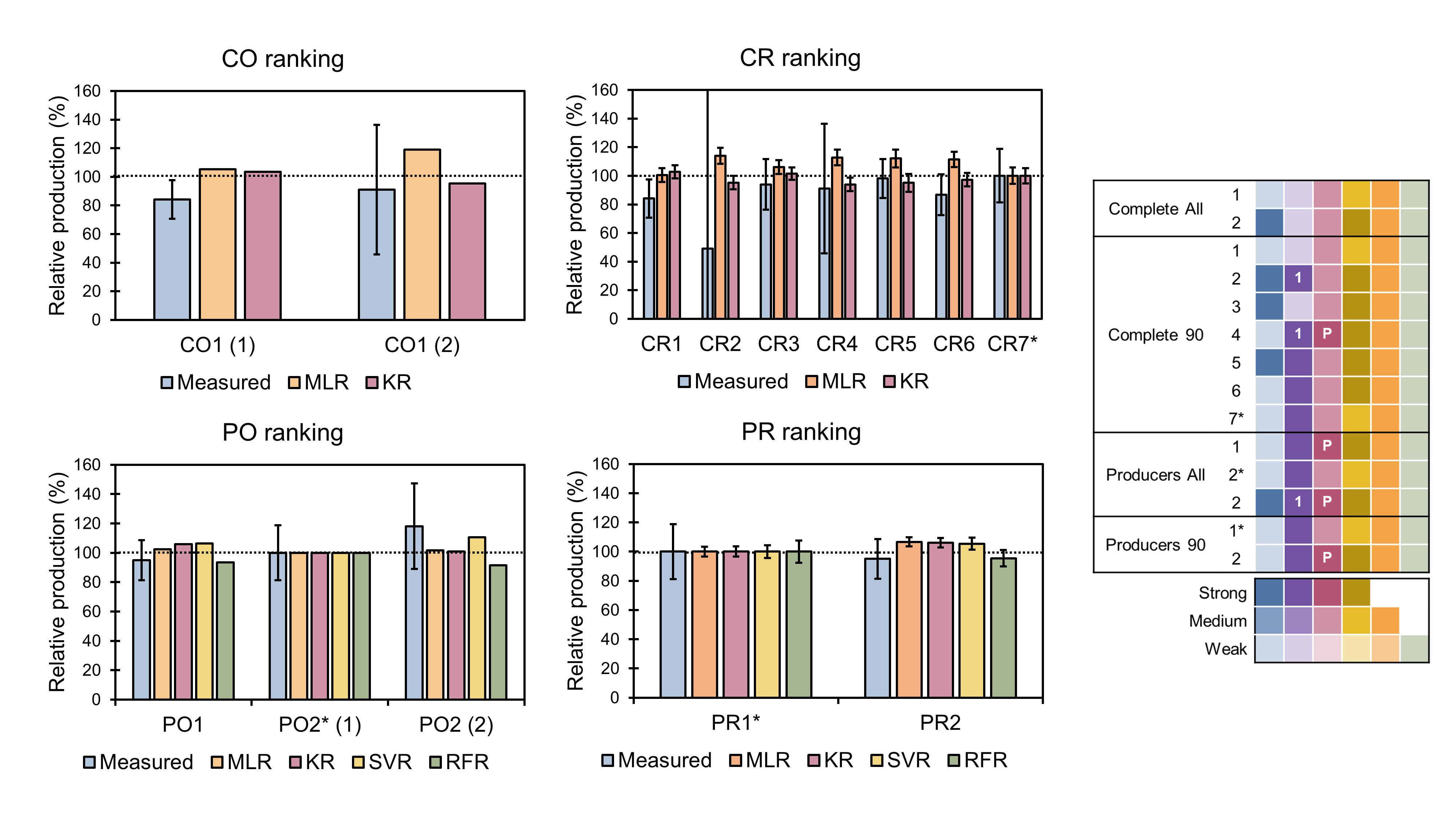

**Figure Sup. 8:** Validation of ML predictions. Comparison of measured and predicted relative production of the predicted top producers relative to the production of the best measured produced (BMP). Each of the left panels represents a different learning strategy . Genotypes of the plotted strains are shown in the right panel where * indicates strains equal to BMP. Factor 1 refers to ARO4, factor 2 refers to AROL expect when 1 is shown (ARO1), factor 3 refers to ARO7 except when P is shown (PHEA), factor 4 refers to PAL; 5 to C4H and 6 to CPR. Promoter strengths are represented by colour intensity. Strain names are defined based on the ranking they belong and their position in the ranking, when two strains share the same position they are followed by (1) and (2). CO, complete dataset with one time training; CR, complete dataset with recurrent training; PR, producers dataset with one time training; PR producers dataset with recurrent training.

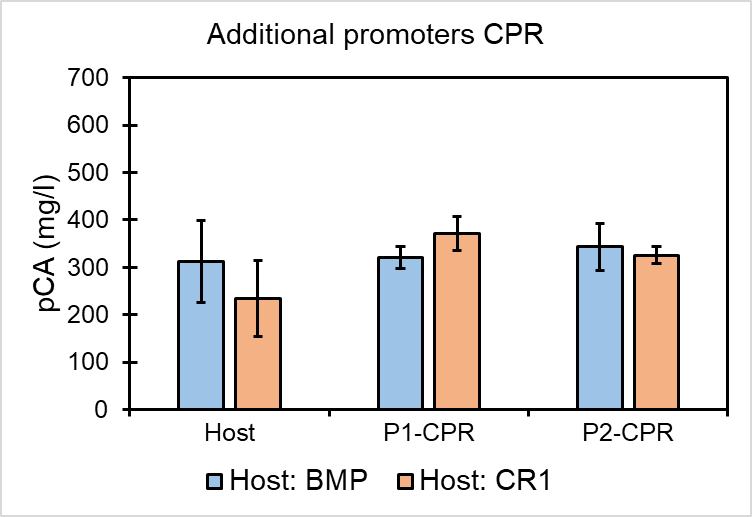

**Figure Sup. 9:** Effect of substituting the CPR promoter in two different hosts: the best measured strain (BMP) and the top producer in the CR ranking (complete dataset recurrent training).

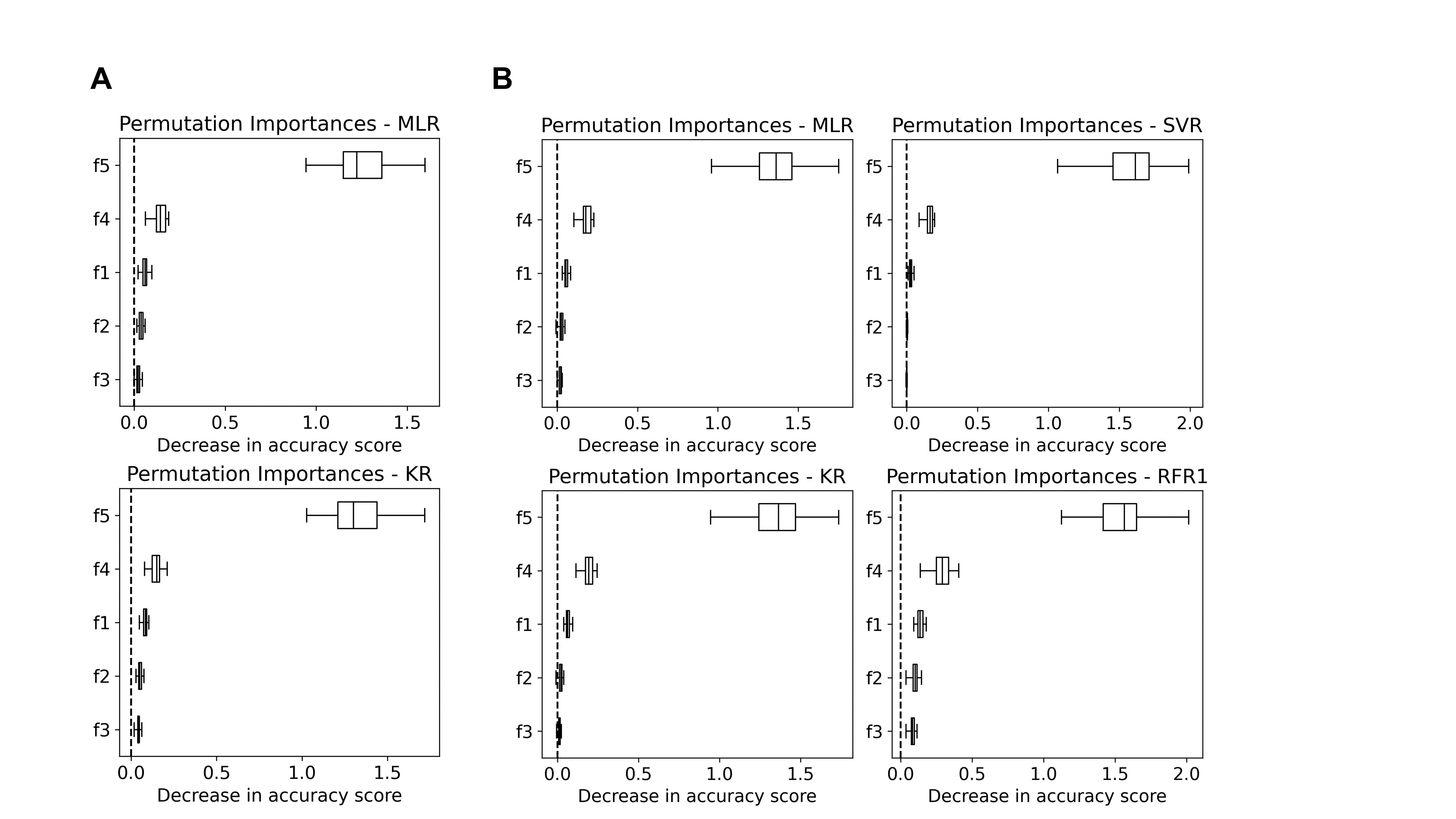

**Figure Sup. 10:** Permutation feature importance results obtained using the *complete* (**A**) or the *producers* (**B**) datasets.

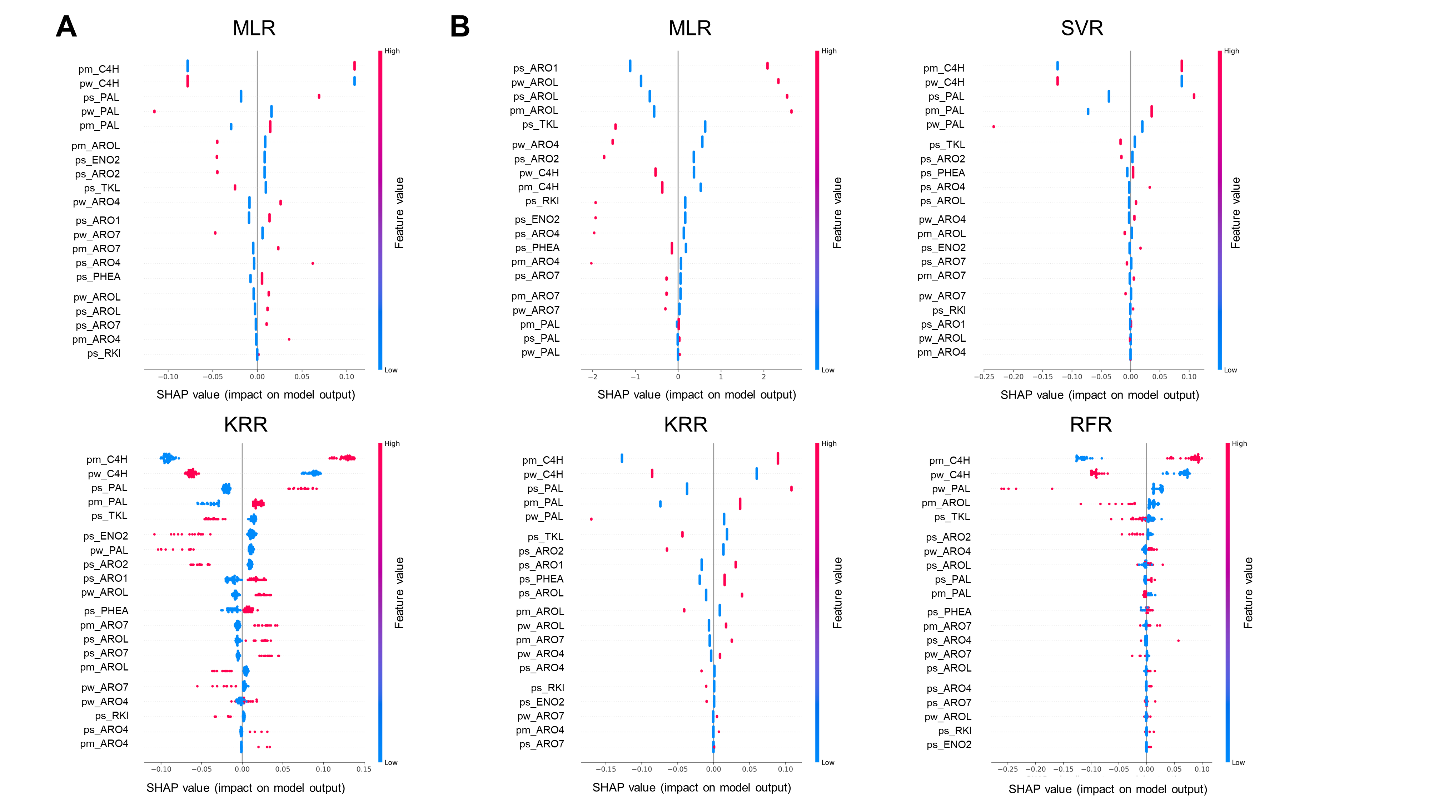
**Figure Sup. 11:** SHAP values obtained using the *complete* (**A**) or the *producers* (**B**) datasets.

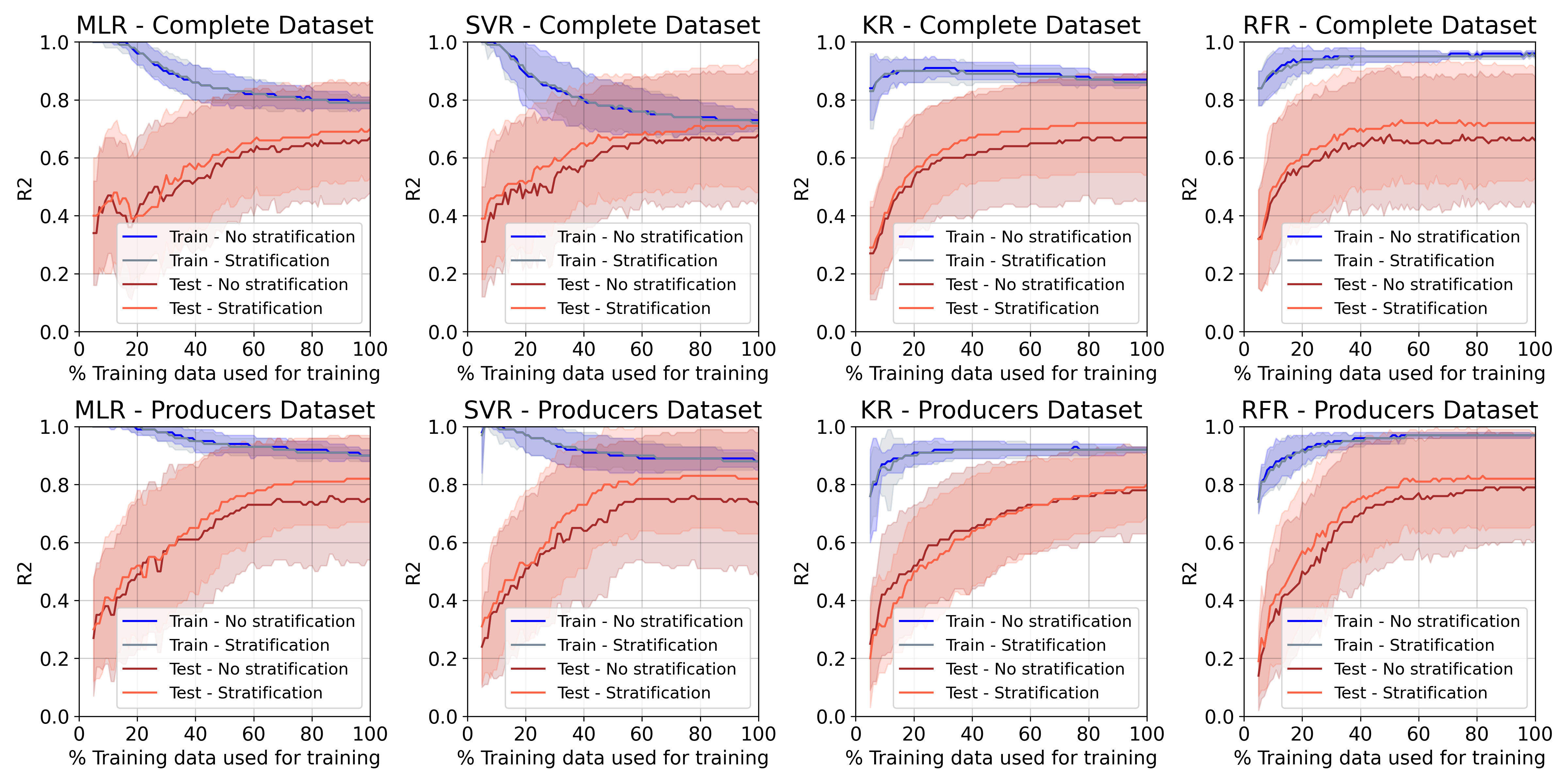

**Figure Sup. 12:** Effect of training data size on accuracy of predictions of different ML algorithms (MLR, multiple linear regression; SVR, support vector regression; KR kernerl ridge regression; RFR, random forest regression) with the complete or producers dataset. Negative R^2^ values obtained for some test-train splits were omitted for the calculation of mean and std (this values are obtained when the average of the training data is a better estimator than the trained model).

**SUPPLEMENTARY REFERENCES**

1. Van Dijken,J.P., Bauer,J., Brambilla,L., Duboc,P., Francois,J.M., Gancedo,C., Giuseppin,M.L.F., Heijnen,J.J., Hoare,M., Lange,H.C., *et al.* (2000) An interlaboratory comparison of physiological and genetic properties of four Saccharomyces cerevisiae strains. *Enzyme and Microbial Technology,* 26, 706-714.

2. Verwaal,R., Buiting-Wiessenhaan,N., Dalhuijsen,S. and Roubos,J.A. (2018) CRISPR/Cpf1 enables fast and simple genome editing of Saccharomyces cerevisiae. *Yeast*, 35, 201-211.
